# A bacterial NLR-related protein senses distinct phage triggers through a single interface

**DOI:** 10.1101/2024.12.17.629029

**Authors:** Laurel K. Robbins, Emily M. Kibby, Amar Deep, Nathan K. Min, Lindsay A. Whalen, Layla D. Borja Najera, Toni A. Nagy, Layla Freeborn, Emily G. Armbruster, Joseph Pogliano, Kevin D. Corbett, Aaron T. Whiteley

## Abstract

Immune systems must rapidly sense viral infections to initiate antiviral signaling, but sensing poses a unique biochemical challenge because viruses rapidly evolve to escape detection. Immune receptors must therefore detect conserved components or activities that are crucial to the viral lifecycle and cannot easily be altered. Here, we show that a bacterial NLR-related protein, bNACHT11, senses viral (phage) infection via direct interactions with multiple phage proteins that are unrelated in their sequences, structures, and functions, thereby limiting viral escape. A conserved surface on the bNACHT11 C-terminal sensor domain binds at least five distinct activators, and a cryo-electron microscopy structure reveals sensing of the protein backbone through β-augmentation. Activator protein binding to bNACHT11 induced oligomerization and effector domain clustering, which limited phage infection by initiating programmed cell death through plasmolysis. These findings reveal a sophisticated immune strategy that counters the rapid evolution of viruses, with parallels to human and plant immune signaling.

## Introduction

Bacteria are under constant threat from bacteriophages (phages) and have evolved sophisticated immune signaling pathways, known as defense systems, to protect themselves. A challenge facing these systems is the sensitive detection of diverse and rapidly evolving phages, each with their own suite of proteins. This selective pressure has driven the evolution of phage-sensing strategies that extend beyond canonical one-to-one interactions between host receptors and phage ligands. For example, some defense systems detect multiple phage proteins through recognition of a conserved protein fold (*1, 2*). Alternatively, some defense systems sense phage protein activities such as perturbations in host processes (*3, 4*), hallmarks of genome destruction (*5–7*), or evidence of immune evasion (*8, 9*). Additional paradigms of phage sensing likely remain to be discovered among the many defense systems whose mechanisms are not yet well understood (*10–14*).

Bacterial NACHT-domain containing proteins (bNACHT proteins) represent a widespread, polymorphic, and poorly understood class of recently identified phage defense systems (*15*). bNACHT proteins are named for their prototypical central NACHT domain, a P-loop NTPase that is a subtype of Signal Transduction ATPases with Numerous Domains (STAND) NTPases (*16*). Proteins containing a NACHT domain (an acronym of the first proteins found to contain this domain) have a unifying role in innate immunity. Many mammalian NACHT containing proteins are nucleotide-binding domain and leucine-rich repeat (NLR) proteins. Human NLRs sense cytosolic signs of infection and contribute to the formation of the inflammasome, a large hetero-oligomeric protein complex which initiates inflammation through cytokine processing and induces programmed cell death (*16*). bNACHT proteins are also related to fungal NLR-related proteins that mediate kin discrimination (*16*) and are more distantly related to plant NLRs, which form oligomeric resistosomes in response to pathogen invasion (*15, 17–20*). A hallmark of NLRs is a conserved domain architecture consisting of a C-terminal sensor domain, a central NACHT/NB-ARC/STAND NTPase domain, and an N-terminal effector domain that coordinates an immune response following activation (**Fig. 1A**).

**Fig. 1.**
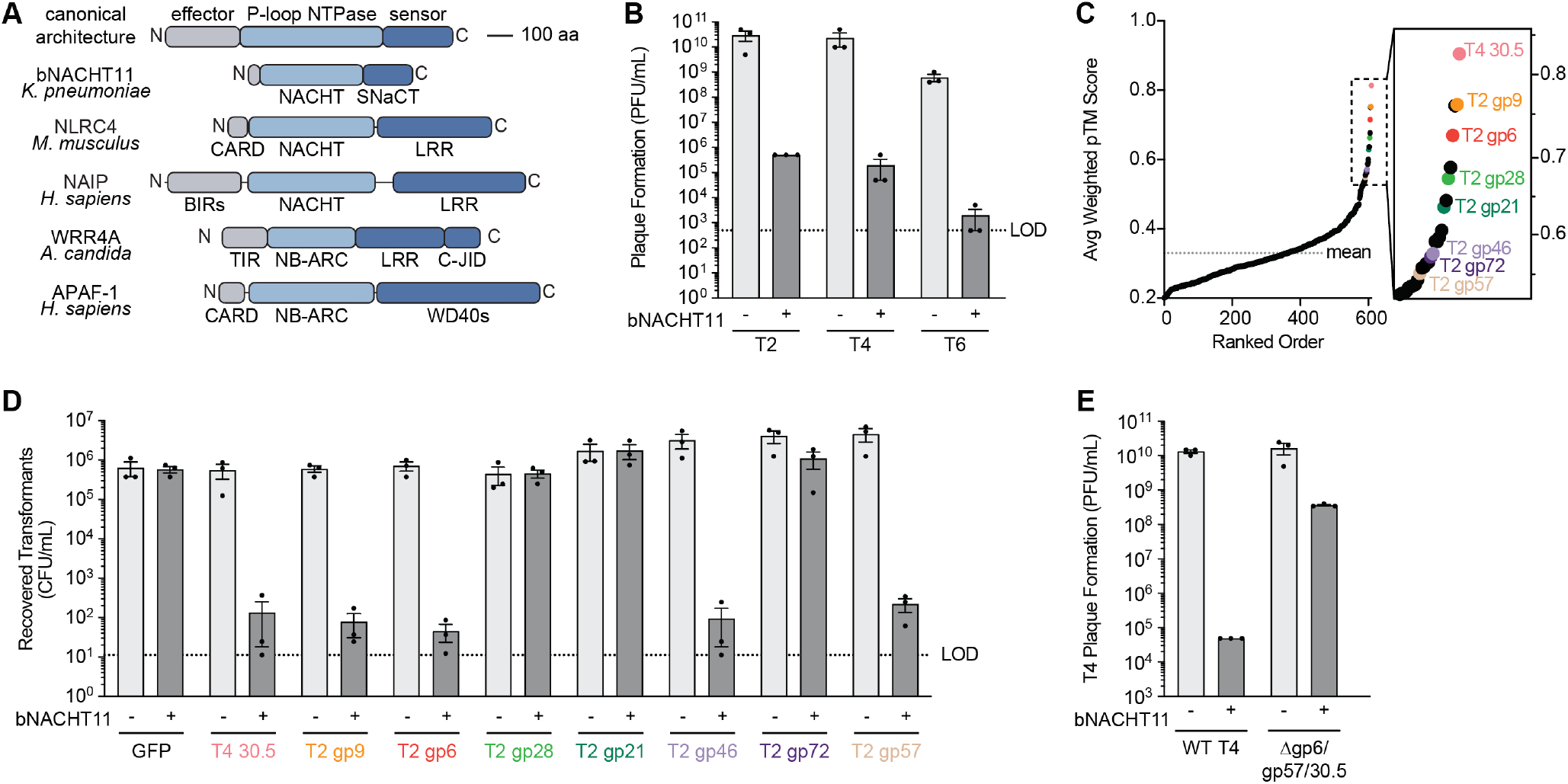
bNACHT11 is activated by multiple phage proteins. **(A)** Domain architecture for bNACHT11, NLRs, and NLR-related proteins highlighting a conserved tripartite domain architecture with an N-terminal effector, central P-loop NTPase, and C-terminal sensor. **(B)** Efficiency of plating of phages T2, T4, and T6 infecting *E. coli* expressing empty vector (-) or bNACHT11 (+). Data represent the mean ± standard error of the mean (SEM) of n = 3 biological replicates, shown as individual points. **(C)** Results of AlphaFold-multimer screening for bNACHT11. Average weighted pTM score (ipTM×0.8 + pTM×0.2) was calculated for each protein screened, then ranked such that the highest-scoring proteins are on the right of the graph (inset). See **Table S2** for the data used to generate this figure. **(D)** Quantification of recovered transformants of *E. coli* expressing an empty vector (-) or bNACHT11 (+) transformed with plasmids encoding GFP or the indicated phage protein. The expression of GFP and phage proteins is IPTG-inducible. Experiments were performed using 500 µM IPTG induction in LB media, and data represent the mean ± SEM of n = 3 biological replicates, shown as individual points. **(E)** Efficiency of plating of wild-type (T4) or mutant (T4 Δgp6/gp57/30.5) phages infecting *E. coli* expressing an empty vector (-) or bNACHT11 (+). Data represent the mean ± SEM of n = 3 biological replicates, shown as individual points.

The most abundant bNACHT proteins encode a conserved domain of unknown function called a Short NACHT Associated C-terminal (SNaCT) domain. SNaCT domains are rapidly evolving and predicted to be essential for phage sensing, but the stimuli for activation is unknown (*15*). Here, we define how a representative of this clade, bNACHT11, detects phage infection and initiates programmed cell death. We show that the SNaCT domain of bNACHT11 binds multiple phage-encoded proteins that differ by their sequence, structure, and function. The SNaCT domain accomplishes this sensing feat by using a binding interface mediated by backbone interactions that are agnostic to amino acid sequence and simultaneous hydrophobic interactions. Activator binding triggers assembly of a heptameric bNACHT11 complex stabilized by interactions between the NACHT domains and adjacent SNaCT domains. Oligomerization of bNACHT11 activates a short, N-terminal effector domain to initiate plasmolysis-mediated cell death, which ultimately limits phage replication to protect the bacterial colony (a process called abortive infection).

## Results

### bNACHT11 is activated by multiple phage proteins

We investigated how the most abundant clade of bNACHT proteins senses phage infection through large-scale, pairwise screening of high-confidence interactions between the immune receptor and phage proteins using AlphaFold-multimer. We selected bNACHT11 as a representative because close homologs are abundant in Gammaproteobacteria, including *Klebsiella* and *Escherichia* species, and because it provides robust defense against phages T2, T4, and T6 (**Fig. 1B**) (*15*). We set up our computational screen by first constructing a non-redundant library of 22,719 open reading frames that represent the diverse pangenome of 91 different *Tequatrovirus* (T4-like) phages (*21, 22*). We clustered proteins by sequence similarity and excluded proteins shorter than 50 amino acids or present in only a single genome to reduce library complexity (**Table S1**, Methods). Next, we used AlphaFold-multimer to perform a high-throughput screen for protein-protein interactions between bNACHT11 and a final set of 629 representative phage proteins. We scored the confidence of each interaction using the weighted pTM score, which incorporates both overall prediction confidence (pTM) and inter-chain prediction confidence (ipTM; **Figs. 1C, S1B-I**). Each interaction was then predicted 5 separate times and the average weighted pTM score was used to score each phage protein. This *in silico* interaction screen identified several phage proteins with high average weighted pTM scores.

We selected proteins encoded by phages T2 and/or T4 with high average weighted pTM scores for experimental analysis and measured their ability to activate bNACHT11 in a genetic assay. bNACHT defense systems defend against phage by initiating cell death (abortive infection) following phage sensing, and hyper-activation of bNACHT11-related proteins inhibits bacterial growth (*2, 13, 15, 23, 24*). Therefore, we constructed a genetic assay by co-expressing phage proteins with bNACHT11 and assessing reduction in colony formation as a proxy for bNACHT11 activation. Co-expression of a non-interacting protein (GFP) with bNACHT11, or phage proteins alone had no effect on colony formation (**Fig. 1D**). However, co-expression of T4 30.5, T2 gp6, T2 gp9, T2 gp46, or T2 gp57 with bNACHT11 resulted in a large reduction in colony formation (**Fig. 1D**).

We next investigated candidate bNACHT11 activators during phage infection by generating single, double, and triple deletions of activators in phage T4. Phage T4 was selected because it encodes the fewest predicted bNACHT11 activators (30.5 and homologs of T2 gp6 and T2 gp57) (**Fig. S2A-F**). All three single deletion mutants partially evaded bNACHT11-mediated defense and deletion of all three activators resulted in near-complete evasion (**Fig. 1E, S2B**). Together, these results suggest that bNACHT11 can detect multiple phage proteins that provide redundant triggers of immune signaling.

### Phage activators trigger bNACHT11 oligomerization

We purified recombinant bNACHT11 and three candidate activators (T2 gp6, T2 gp9, and T2 gp57) and measured their binding affinity using microscale thermophoresis. We found that bNACHT11 binds gp6 with a dissociation constant (*K*_*D*_) of 80 ± 21 nM, T2 gp9 with a *K*_*D*_ of 269 ± 43 nM, and T2 gp57 with a *K*_*D*_ of 149 ± 68 nM (**Figs. 2A-C, S3I–L**). Next, we determined if phage protein binding induces bNACHT11 oligomerization. Ligand binding by human and plant NLRs induces a conformational change in the core NTPase that promotes higher-order oligomerization (*25, 26*). This transition is typically dependent on nucleotide exchange; P-loop NTPases generally bind a nucleoside diphosphate (NDP; ADP or GDP) in their inactive/monomeric state and a nucleoside triphosphate (NTP;ATP or GTP) in their activated/oligomeric state (*16*). We measured bNACHT11 oligomerization following activator binding using size-exclusion chromatography. Incubation of bNACHT11 with ATP and T2 gp6, T2 gp9, or T2 gp57 resulted in the formation of a higher-order oligomeric complex (**Fig. 2D-F**). Oligomerization required ATP, but not ATP hydrolysis, as incubation of bNACHT11 with the nonhydrolyzable ATP analog AMP-PNP and T2 gp57 still resulted in complex formation (**Fig. 2F**). Incubation of bNACHT11, T2 gp6, and GTP did not result in complex formation (**Fig. S3M**). These findings suggest that phage activators trigger nucleotide-dependent oligomerization of bNACHT11 similarly to ligand-dependent oligomerization of inflammasome components in humans or resistosome assemblies in plants (*26, 27*).

**Fig. 2.**
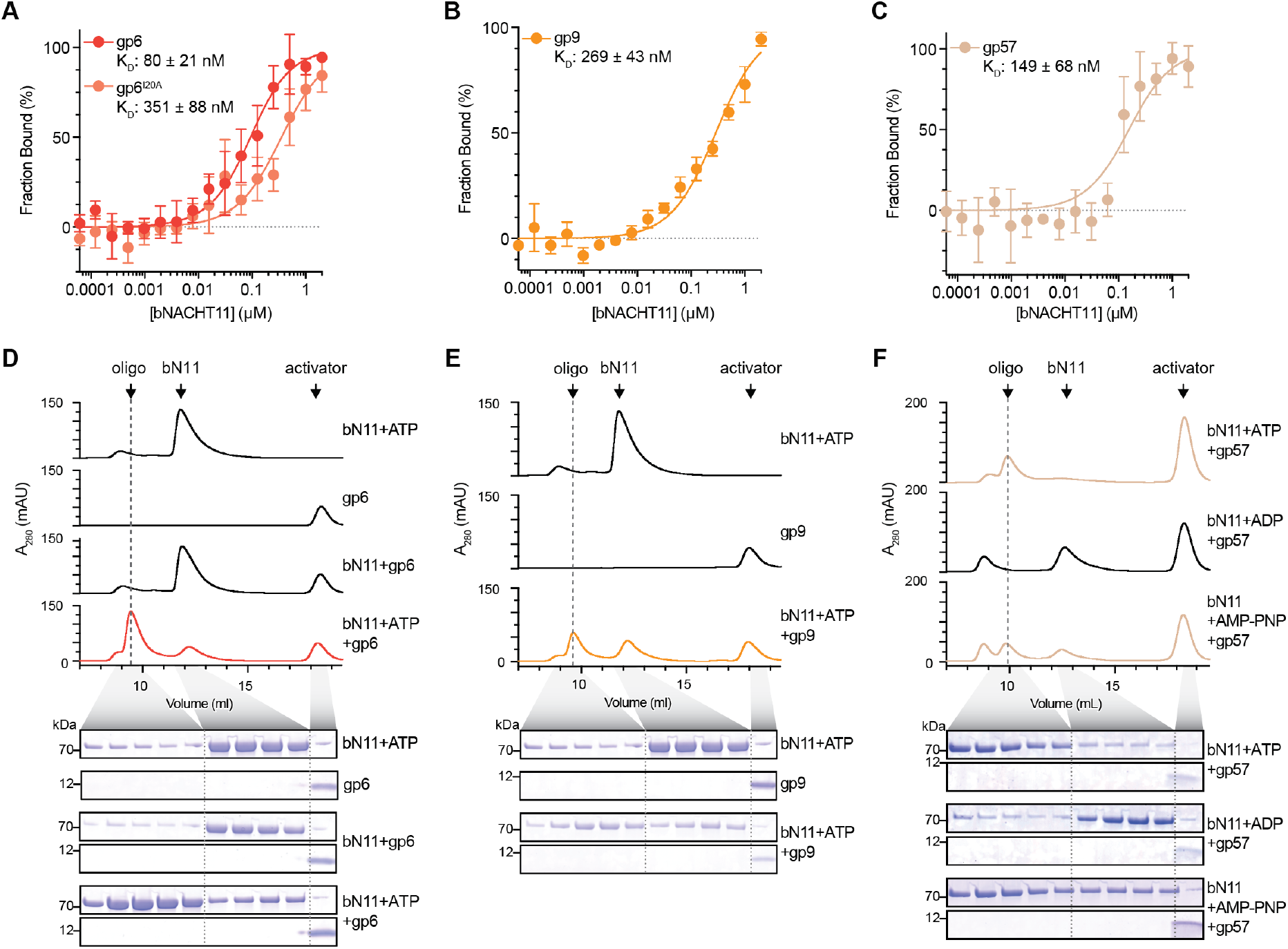
Direct binding of phage protein activators to bNACHT11 induces oligomerization. **(A-C)** Binding curve of purified gp6 and gp6^I20A^ (A), gp9 (B), or gp57 (C) binding to purified bNACHT11, measured using microscale thermophoresis (MST). Data in A represent the average of n = 4 replicates ± SEM. Data in B and C represent the average of n = 3 replicates ± SEM. See **Figure S3** for protein purification gels, untransformed individual replicates, and binding measurements in high salt conditions. **(D-F)** Above: Traces showing the absorbance at 280 nm (A_280_) of the indicated samples run over a Superdex 200 Increase size exclusion (SEC) column. Labels indicate the identity of proteins in each peak. Below: SDS-PAGE analysis of representative samples taken from collected fractions in indicated volumes. For samples containing both bNACHT11 and an activator, portions of the gel corresponding to each are shown. Data in (D) are representative of n = 3 replicates. Data in (E) are representative of n = 2 replicates, traces and gels for bN11+ATP are reused from (E). Data in (F) are representative of n = 2 replicates.

### Structure of phage activator–bound bNACHT11 complex

To understand the molecular basis of infection sensing and the overall architecture of activated bNACHT11, we determined a ∼2.5 Å-resolution cryo-electron microscopy (cryoEM) structure of bNACHT11 in complex with T2 gp57 (**Fig. 3A-B, S4, Table S5**). The structure reveals that bNACHT11 forms a wheel-like assembly, in which the C-terminal SNaCT domain of each protomer engages T2 gp57, forming a 7:7 stoichiometric complex (**Fig. 3A**). The NACHT and SNaCT domains have an ordered radial arrangement while the N-terminal 45 residues expected to be the effector domain did not resolve, likely due to conformational flexibility. The heptameric complex measures approximately 18 × 18 × 11 nm (**Fig. 3B**) and resembles the activated complex of other NLR-related proteins (*28–30*). An analysis of structures similar to the bNACHT11 protomer identifies the top hits as eukaryotic NLRs, including NLRC4 from *Mus musculus* (PDB: 3JBL) and NAIP from *Homo sapiens* (PDB: 8FVU) (**Fig. S5A**). A comparison of the NACHT domains across these hits highlights the striking structural similarity between bNACHT11 and mammalian NLR proteins (**Fig. S5B**)

**Fig. 3.**
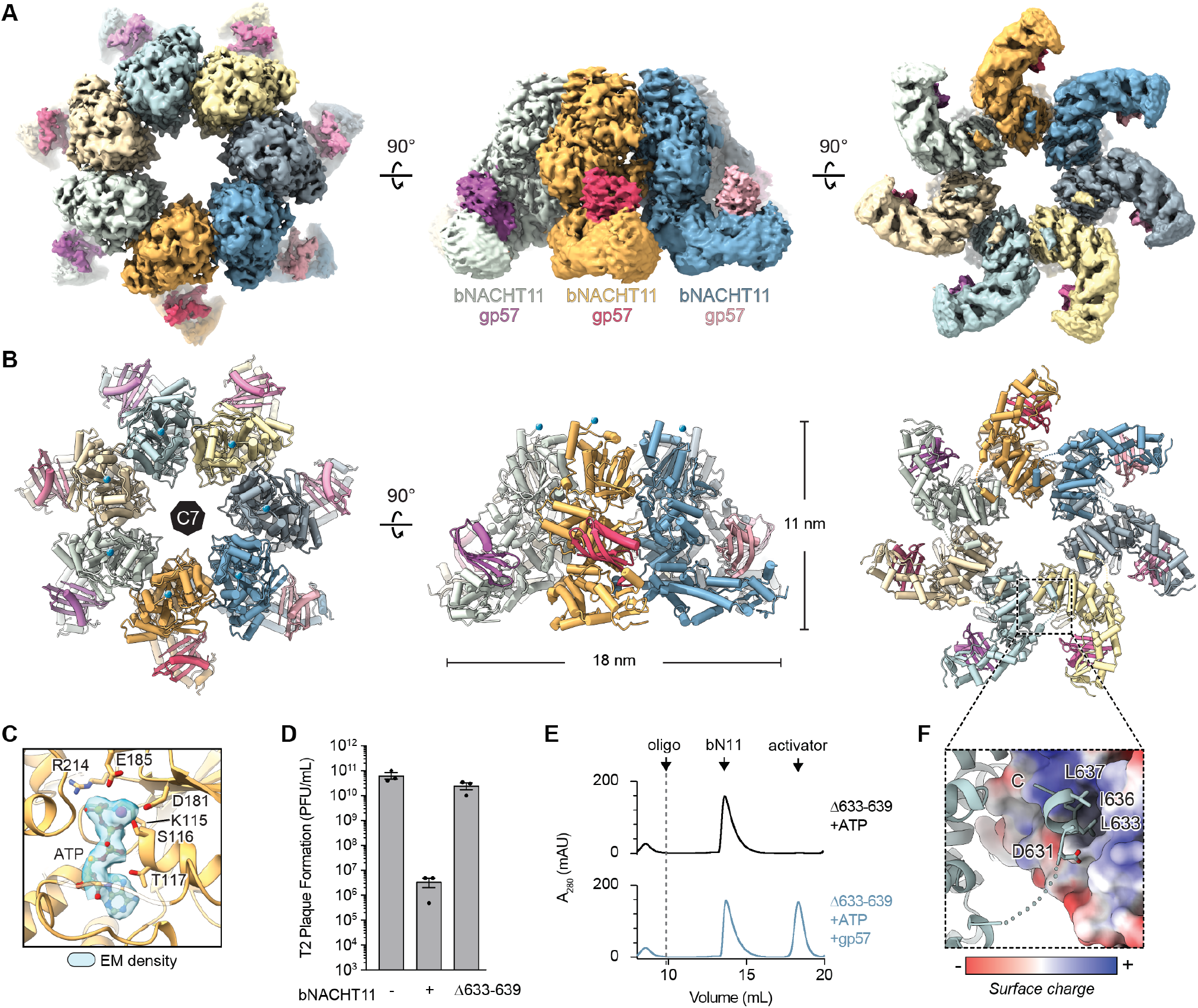
CryoEM structure of bNACHT11–gp57 complex. **(A)** Three views of the cryoEM density map of bNACHT11 in complex with T2 gp57 showing a heptameric activated state. **(B)** Cartoon view of views as in (A) of the bNACHT11–gp57 complex. **(C)** EM density of modeled ATP and Mg^2+^ ions in the nucleotide binding pocket of the bNACHT11 NACHT domain. Side-chain for residues predicted to interact with the ligand are shown as sticks. **(D)** Efficiency of plating of phage T2 infecting *E. coli* expressing an empty vector (-), bNACHT11-6×His, or bNACHT11-6×His with a deletion of residues 633-639 (Δ633-639). Data represent the mean ± SEM of n = 3 biological replicates, shown as individual points. **(E)** Traces showing the absorbance at 280 nm (A_280_) of the indicated samples run over a size exclusion column. Labels indicate the identity of proteins in each peak. **(F)** Zoomed-in view of the C-terminal tail docking into the SNaCT pocket of a neighboring protomer. The pocket surface is colored by electrostatic potential, where white indicates neutral, blue indicates positive, and red indicates negative charge.

The structure of activated bNACHT11 also allows us to interrogate how the NACHT and SNaCT domains cooperate to assemble the active complex. The primary oligomerization interface is composed mainly of charged residues in the NACHT domain (**Fig. S5C**). We designed five different charge-swap mutations to test the biological relevance of this interface by measuring their ability to protect against phage T2 (**Fig. S5D**). Four of these mutants expressed at wild-type levels and each abolished phage defense, supporting the importance of this oligomerization interface. A careful examination of this structure also revealed that the extreme C-terminus of bNACHT11 (residues 633-639) extends into a pocket within the SNaCT domain of a neighboring protomer (**Fig. 3F**). We hypothesized this interaction may stabilize the overall wheel-like architecture of the activated complex and generated a mutant with a deletion of the C-terminal tail (Δ633-639). bNACHT11 Δ633-639 was unable to form an oligomeric complex following incubation with ATP and T2 gp57 (**Fig. 3E**). Additionally, this mutant did not provide phage defense (**Fig. 3D**).

Our structure resolved ATP·Mg^2+^ bound within the NACHT domain at the expected location. ATP·Mg^2+^ interacts with conserved residues in the Walker A motif (K115) and a previously predicted ATP binding residue (R214) (**Fig. 3C**) (*15*). This finding is consistent with the requirement of ATP for oligomerization of bNACHT11 and other NLRs (**Fig. 2 D-F**) (*16*).

### bNACHT11 binds phage activators through β-augmentation and hydrophobic interactions

Surprisingly, T2 gp6, T2 gp9, and T2 gp57 are not related by sequence or predicted structure (**Fig. S3A**). Yet AlphaFold models predicted that bNACHT11 binds to all three activators using the same interface of the SNaCT domain and our structure of bNACHT11–T2 gp57 supports that model. Other defense systems recognize multiple ligands that share a conserved structure, or use overlapping but distinct binding sites for different activators (*31*). bNACHT11 is different because it recognizes structurally diverse phage proteins using the same binding interface (**Figs. 4A-C**).

**Fig. 4.**
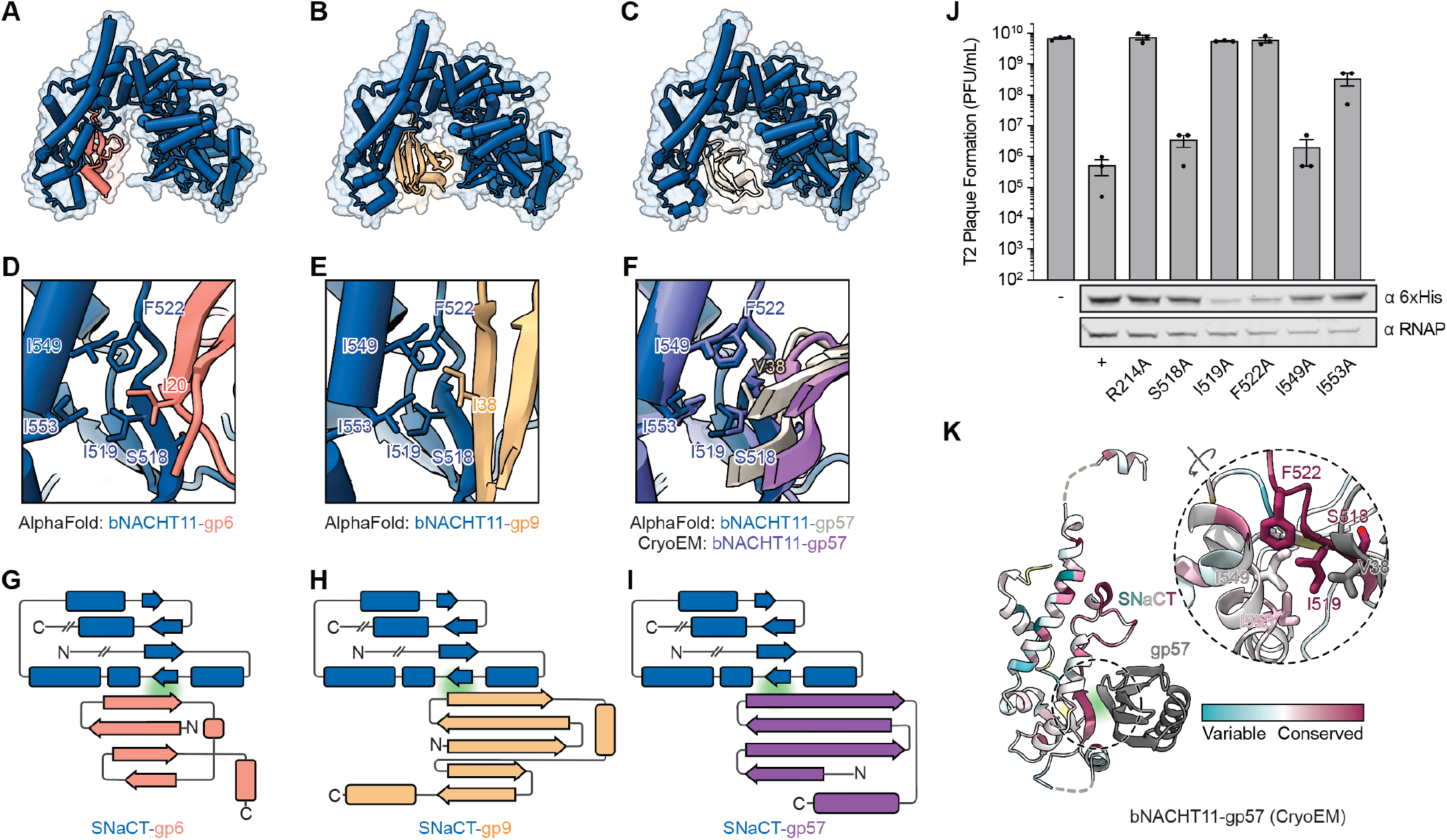
bNACHT11 activation is mediated by hydrophobic contacts and β-augmentation. **(A-C)** AlphaFold-multimer predicted structure of bNACHT11 (blue) binding T2 gp6 (red) (A), T2 gp9 (orange) (B), and T2 gp57 (tan) (C). (**D-F**) Zoomed in view of the AlphaFold-multimer predicted binding interface between bNACHT11 (blue) and the indicated phage protein. Residues predicted to be involved in the interaction are labeled and shown as stick models. For T2 gp57, the cryoEM structure is overlayed in purple (F). **(G-I)** Topology diagrams manually drawn from the cryoEM structure of SNaCT domain and AlphaFold predictions of T2 gp6 and T2 gp9 (G-H) and the cryoEM structure of T2 gp57 (I). The blue arrowing corresponding to bNACHT11, highlighted with green, is not to scale and is drawn to indicate the involved β-augmentation interaction with bNACHT11. **(J)** Top: Efficiency of plating of phage T2 infecting *E. coli* expressing an empty vector (-) or the indicated genotype of bNACHT11-6×His. Data represent the mean ± SEM of n = 3 biological replicates, shown as individual points. Bottom: Western blot analysis of *E. coli* expressing 6×His-tagged bNACHT11 of the indicated genotype. Representative image of n = 2 biological replicates. (**K**) ConSurf-mapped conservation of the SNaCT domain interacting with T2 gp57. The SNaCT domain is colored by conservation as indicated.

We investigated the molecular details for how the SNaCT domain binds different phage ligands by comparing our structure of bNACHT11–T2 gp57 and predicted complexes of bNACHT11 and T2 gp6 or T2 gp9. The first pattern of interaction we observed is that the edge of a β-strand of each activator made backbone interactions with a β-strand of the SNaCT domain of bNACHT11, forming an anti-parallel β-sheet (**Fig. 4G-I**), a protein-protein interaction called β-augmentation. β-augmentation was observed for all three activators where the same strand of bNACHT11 interacts with an outside strand from the phage protein activator (**Fig. 4G-I**). Notably, the interacting strands of the phage protein activators share no sequence identity: IAI for gp6, DIA for gp9, and GVV for gp57. These findings are in line with β-augmentation because the hydrogen binding that completes the newly formed β-sheet is mediated by the peptide backbone and agnostic to amino acid side chain.

The second pattern of interaction we observe is that all three phage proteins interact with the SNaCT domain through a hydrophobic contact. We generated alanine substitutions at this interface (bNACHT11 S518, I519, F522, I549, and I553) and within the nucleotide binding domain (bNACHT11 R214) as a control (**Fig. 4J**). Mutations I519A and F522A decreased bNACHT11 expression, precluding further analysis (**Fig. 4J**). In contrast, variants I553A, S518A, and I549A were expressed at wild-type levels, and all three reduced phage defense (**Fig. 4J**). An analysis of amino acid conservation for the SNaCT domain shows that, although the overall SNaCT domain has high sequence diversity, many of the hydrophobic residues along this short β-strand region are conserved, consistent with a crucial role in activator binding (**Fig. 4K, S6A**).

To further validate the predicted binding interface between bNACHT11 and T2 gp6, we mutated I20 of T2 gp6 to alanine. This residue is predicted to interact with the hydrophobic interface of bNACHT11. We found that the gp6 I20A mutation attenuated growth inhibition when co-expressed with bNACHT11 (**Fig. S3N**). Consistently, purified T2 gp6 I20A exhibited reduced affinity for bNACHT11, with a *K*_*D*_ of 351 ± 88 nM (**Figs. 2A, S2J**). To assess the contribution of ionic interactions *in vitro*, we performed binding experiments between bNACHT11 and T2 gp6 under high-salt conditions (1 M KCl) and found that the binding affinity was largely unaffected (**Fig. S3G–H)**. Together, these results support that β-augmentation and hydrophobic interactions at the SNaCT-domain interface of bNACHT11 are critical for activator recognition.

### bNACHT11 activators bind distinct host proteins

Our results demonstrate that bNACHT11 recognizes multiple unrelated phage proteins, however, the functions of these proteins are unknown. We investigated whether these proteins interact with host factors by generating a functional epitope-tagged version of each activator, expressing each protein in *E. coli* in the absence of bNACHT11, and performing an immunoprecipitation. Samples were silver-stained and gels were compared between epitope tagged and untagged controls. T2 gp6, T4 30.5, and T2 gp57 co-purified with distinct host proteins (**Figs. 5A, 5D, 5G**). Mass spectrometry identified the T2 gp6 interactor as pantothenate kinase (CoaA) (**Fig. 5B**), the T4 30.5 interactor as polyphosphate kinase (Ppk) (**Fig. 5E**), and the T2 gp57 interactor as glycerol kinase (GlpK) (**Fig. 5H**). We validated these interactions by co-expressing VSV-G tagged activators with FLAG tagged host protein and performed reciprocal immunoprecipitation using the FLAG tag (**Figs. 5C, 5F, 5I**). This analysis confirmed the interactions identified by mass spectrometry. Each of these host proteins have well-characterized enzymatic activities and phosphorylate distinct substrates in the cell (**Figs. 5J, 5K, 5L**) (*32–34*). The finding that each bNACHT11 activator binds different host kinases suggests these proteins are functionally distinct.

**Fig. 5.**
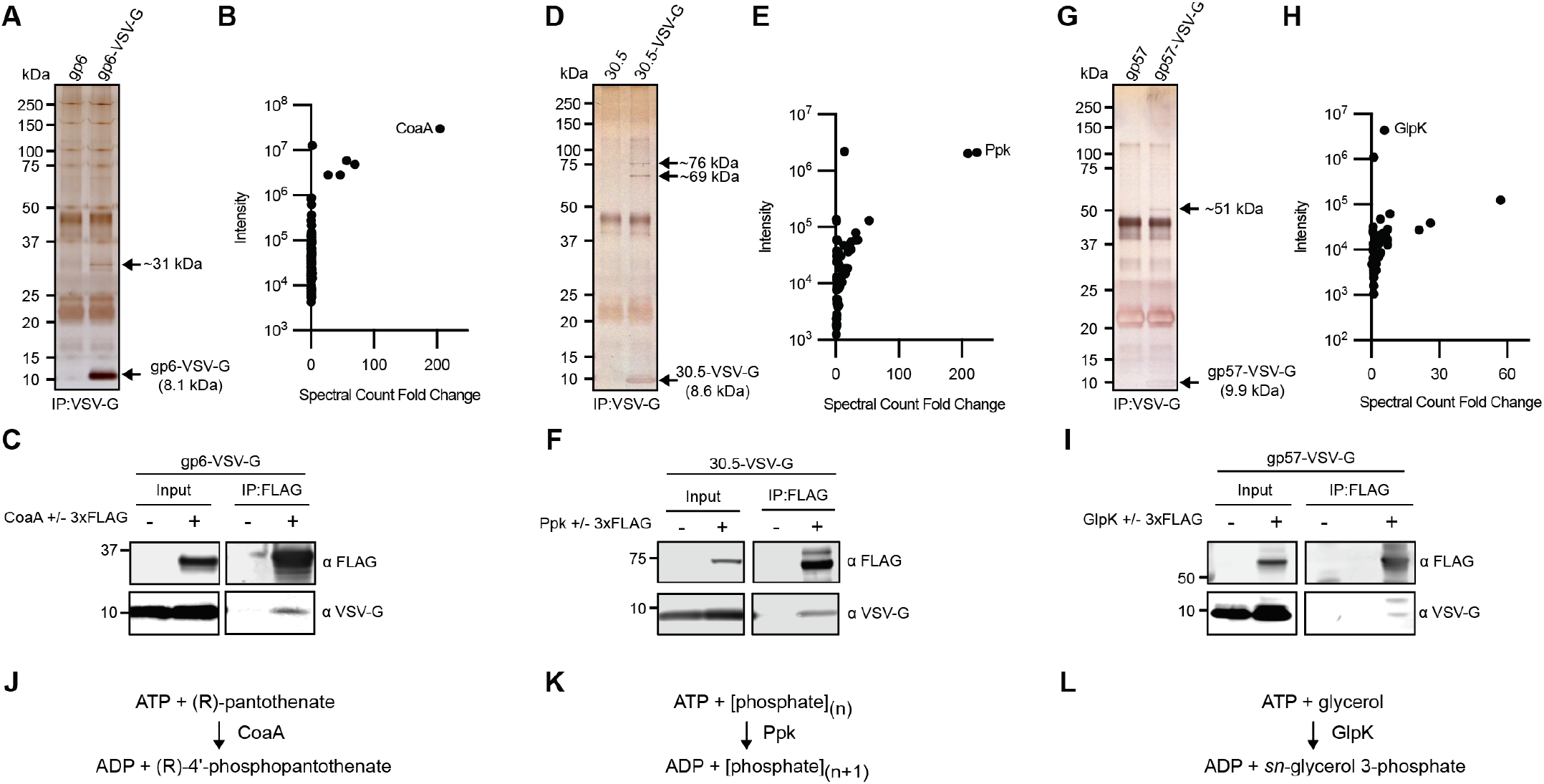
bNACHT11 activators interact with distinct host proteins. **(A, D, G)** Silver-stained SDS-PAGE gels of untagged vs. VSV-G tagged samples following immunoprecipitation (IP) of VSV-G. **(B, E, H)** Mass spectrometry analysis of excised bands from (A), (B), and (C), respectively. Data are plotted as spectral count fold change between tagged and untagged samples vs intensity of the tagged sample. **(C, F, I)** Western blot analysis of co-immunoprecipitations between VSV-G phage protein and FLAG tagged vs. untagged hit identified by mass spectrometry. Input samples represent clarified lysate samples prior to IP by FLAG. **(J, K, L)** Reactions catalyzed by host kinase (*32–34*).

### Activation of bNACHT11 induces plasmolysis

We next determined how activation of bNACHT11 leads to programmed cell death. NACHT domain-containing proteins typically trigger programmed cell death through an N-terminal effector domain that becomes clustered following oligomerization (*15*). bNACHT11 lacks a readily identifiable effector domain and instead contains a short ∼45 amino acid N-terminal extension. This region was not resolved in the cryoEM structure of activated bNACHT11 (**Fig. 3A-B**). To determine if residues within this N-terminal extension were required for phage defense, we conducted mutagenesis of this region (**Fig. 6A, S7A-B**). Additions and substitutions at the N-terminus abolished phage protection, including alanine substitution of two highly conserved residues in this sequence: bNACHT11 D2A and P3A (**Fig. 6A-B, S7A-B**). We were unable to generate deletions within the N-terminus without affecting expression levels (**Fig. S7A**).

**Fig. 6.**
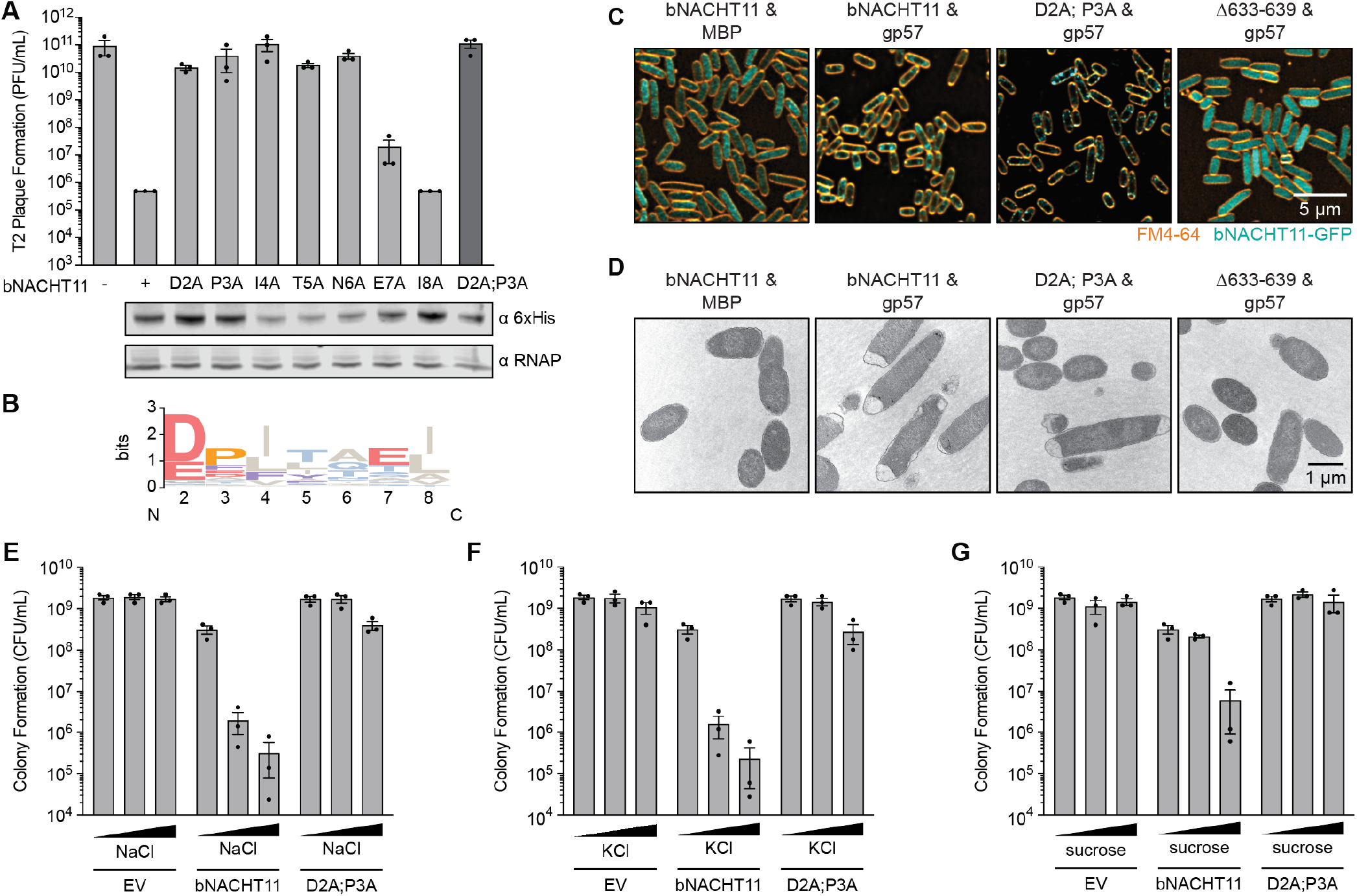
bNACHT11 N-terminal extension induces plasmolysis upon activation. **(A)** Top: Efficiency of plating of phage T2 infecting *E. coli* expressing an empty vector (-) or the indicated genotype of bNACHT11-6×His. Data represent the mean ± SEM of n = 3 biological replicates, shown as individual points. Bottom: Western blot analysis of *E. coli* expressing 6×His-tagged bNACHT11 of the indicated genotype. Representative image of n = 2 biological replicates. **(B)** Sequence logo for the first 8 N-terminal residues of 537 bNACHT sequences from clade 14 (excluding Met encoded by the start codon) (*15*). Data are depicted as bits and represented by the height of each residue. **(C)** Fluorescence laser scanning confocal microscopy of *E. coli* expressing the indicated bNACHT11-GFP genotype with either maltose binding protein (MBP) or T2 gp57. Membranes stained with FM4-64 (orange). Representative of n = 2 biological replicates. **(D)** Transmission electron microscopy (TEM) imaging of the strains shown in A. **(E-G)** Quantification of colony formation of *E. coli* expressing an empty vector (EV), bNACHT11, or D2A; P3A, coexpressed with T2 gp9. The expression of gp9 is IPTG-inducible.Experiments were performed using 500 µM IPTG induction in LB media lacking salt with the addition of (E) no NaCl (0 µM), medium NaCl (91 mM, conventional LB), high NaCl (300 mM), (F) no KCl (0 µM), medium KCl (91 mM), high KCl (300 mM), or (G) no sucrose (0 µM), medium sucrose (91 mM), high sucrose (300 mM). Data represent the mean ± SEM of n = 3 biological replicates, shown as individual points.

We hypothesized that this N-terminal extension functions as the effector responsible for bNACHT11-mediated cell death. This small motif was conserved with a consensus sequence of “MDPIT” and is grossly similar in residue properties to the “MEPIS”-like motifs found in fungal domains and two other antiphage systems that mediate cell death (*35–37*). To interrogate the mechanism of programmed cell death, we used confocal fluorescence microscopy of bacteria expressing GFP-tagged bNACHT11 to visualize changes in localization following activation. We were unable to construct a functional GFP fusion at either terminus (data not shown), however, we successfully engineered an internal GFP-tagged bNACHT11 that remained functional (**Fig. S7C**). Fluorescence microscopy revealed that when bNACHT11 is co-expressed with a non-activating protein control, the bNACHT11 signal was diffuse throughout the cell **(Fig. 6C)**. However, co-expression with T2 gp57 for 30 minutes resulted in distinct puncta near the cell membrane (**Fig. 6C**). The D2A;P3A double mutant of bNACHT11 also formed puncta upon activation despite diminished inhibition of growth when co-expressed with T2 gp57 (**Fig. 6C and S7C**). In contrast, the oligomerization-disrupting Δ633-639 mutant failed to form puncta (**Fig. 6C**). These data suggest that bNACHT11 may localize to the membrane following activation and, while the D2A; P3A mutation did not disrupt localization or puncta formation, these residues are crucial for later stages of signaling.

We investigated the cellular consequences of bNACHT11 activation using transmission electron microscopy to examine cellular morphology. Cells expressing bNACHT11 and T2 gp57 appeared to have collapsed inner membranes near the pole regions (**Fig. 6D**). This phenotype was less common in strains co-expressing T2 gp57 with the D2A; P3A mutant and was absent in the bNACHT11 Δ633-639 mutant. The phenotype appeared similar to literature reports of cells with activated antiphage systems encoding “MEPIS”-like motifs (*35*).

We recognized bNACHT11-mediated cell death as visually similar to plasmolysis, a process that occurs under high osmotic stress, where the inner membrane collapses due to outward osmotic flow of water (*38–40*). We hypothesized that bNACHT11 may induce plasmolysis through interfering with cellular solute homeostasis, for example, by changing the ability of cells to regulate internal ion concentrations and turgor pressure. To test this hypothesis, we compared co-expression of bNACHT11 with the activator T2 gp9 in LB media with varying concentrations of NaCl, KCl, and sucrose. Strikingly, in the absence of additional salt, activation of bNACHT11 no longer inhibited colony formation (**Fig. 6E-G**). Increasing osmotic stress with added salt or sucrose increased growth inhibition following bNACHT11 activation (**Fig. 6E-G**). These observations were specific to bNACHT11 activation because they were not observed for T2 gp9 in the absence of bNACHT11 or bNACHT11 in the absence of activator (**Fig. 6E-G, S7D-F**). Additionally, solute dependent toxicity was almost completely abolished when the D2A; P3A mutant was co-expressed with T2 gp9 (**Fig. 6E-G**). Together, these data show that bNACHT11 activation induces cell death by triggering plasmolysis.

## Discussion

Here, we show that the phage defense system bNACHT11 senses infection by directly detecting multiple unrelated phage proteins using a common interface (**Fig. 7**). This mechanism of phage sensing has not been previously observed but is an ideal immune strategy in the endless competition between bacteria and phages. While CapRel^SJ46^ and KpAvs2 defense systems are also activated by more than one phage protein, these sensors use distinct binding sites for each ligand. Similarly, Avs4 detects phage portal proteins sharing ≤10% amino acid identity by taking advantage of their conserved protein fold. Our analysis reveals that bNACHT11 is capable of binding many unrelated phage proteins by using at least two molecular features. First, β-augmentation between antiparallel β-strands in bNACHT11 and the phage activator contribute to this interaction. These main-chain hydrogen bonds likely provide for specificity and positioning of the phage activator, while limiting phage escape because most sidechain mutations would have a limited impact on these intermolecular interactions. Second, hydrophobic interactions between bNACHT11 and phage activators likely provide the high affinity of the interaction. A possible third requirement is that activators may need to be small enough to fit into the binding pocket, however, we are unable to test this.

**Fig. 7.**
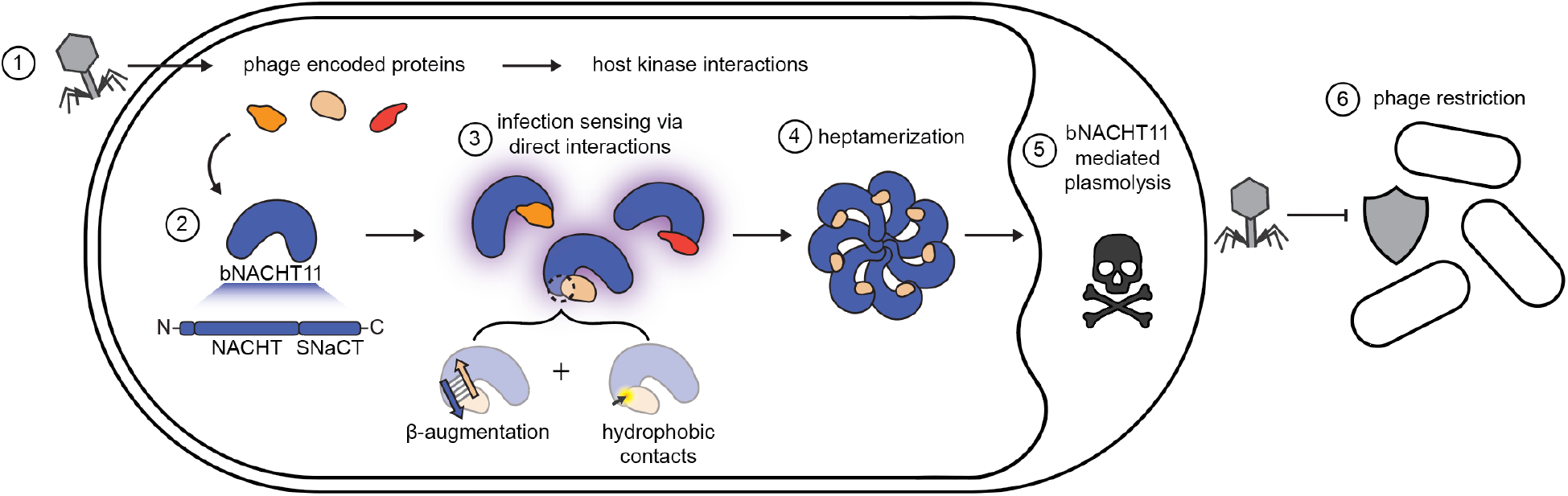
Proposed model for phage protection by bNACHT11. Proposed phage sensing and protection mechanism of bNACHT11. Phage encoded proteins including T2 gp6, T2 gp9, T2 gp57, and T4 30.5 activate bNACHT11 through direct interactions with the SNaCT domain, with interactions mediated by β-augmentation and hydrophobic contacts. Upon activation, bNACHT11 heptamerizes and causes programmed cell death via induction of plasmolysis.

Our findings raise the question of why phages retain bNACHT11 activating proteins when they trigger a potent immune response. We hypothesize that the structure of T2 gp57 and other phage activators is required for their function and that their function provides a selective benefit to phage replication. Our data shows that each phage protein interacts with a core metabolic enzyme, therefore, these proteins may be required to reprogram metabolism and optimize phage replication. Deletions of phage activators did not affect titer or plaque morphology in the absence of bNACHT11, suggesting that these proteins are not essential under the experimental conditions tested. However, the hypothesis that these proteins are conditionally essential is consistent with the mosaic distribution of T2 *gp6*, T2 *gp9*, and T2 *gp57* in the pangenome of *Tequatrovirus* genus, at 95%, 57%, and 91% of the genomes surveyed, respectively (**Table S1**). It is well known that phages reprogram their hosts in numerous ways and our data support previous reports showing phages modulate metabolism to redirect resources to support phage replication (*41*). bNACHT11 acts as an ideal innate immune sensor because it detects redundant phage triggers. The superior function of the SNaCT domain likely underlies why this clade of bNACHT proteins, clade 14, is so abundant in bacteria genomes.

Identification of bNACHT11 activators using AlphaFold-multimer highlights the power of *in silico* strategies for discovering protein-protein interactions (*42*). As AlphaFold becomes increasingly used for identifying protein-protein interactions (*43*), inhibitors (*44–46*), and more across the tree of life (*47, 48*), it is increasingly important to understand opportunities and limitations of this strategy.

The observation that heptamerization of bNACHT11 is stabilized by interactions between NACHT domains and between neighboring SNaCT domains reveals how both domains cooperate to assemble the active complex. This observation reveals an emerging theme in NLR signaling as a similar observations were recently made for the plant NLRs WRR4A-CCG40N and CC_G10_-NLR (*28, 49*). Heptamerization is required for initiation of programmed cell death via the N-terminal extension, however, it is unclear how this region of the protein exactly orchestrates plasmolysis. Ibarlosa et al. found that the N-terminal residues of bNACHT11 (MDPIT) share sequence similarity with two other antiphage systems and a fungal NLR-related protein (consensus motif of “MEPIS”), and that these systems result in phenotypes similar to what we observe (*35*). Our data complement these findings and suggest MDPIT/MEPIS-mediated phenotypes are the result of disrupted osmoregulation and plasmolysis. These data further show that convergent mechanisms of clustering MDPIT/MEPIS motifs initiate programmed cell death in diverse organisms. Given the distance between *E. coli* and filamentous fungi, we hypothesize that these domains may interact with membranes directly and not via an intermediate protein. However, it is unclear how membranes might be perturbed biophysically because the N-terminus of bNACHT11 is not obviously enriched for hydrophobic residues.

Our results reveal a complete activation pathway that links phage sensing to effector-mediated cell death and abortive infection for a bacterial NLR-related protein. The prevalence of NACHT and related STAND NTPase domain containing proteins across the tree of life is not random. Instead, these protein domains appear uniquely adapted to accurately and sensitively make the high stakes decision to induce programmed cell death. bNACHT11 has further evolved the extraordinary ability to recognize a broad range of phage proteins. It remains to be discovered if other immune sensors, including those in humans, may also use β-augmentation to increase the breadth of their interaction.

## Acknowledgements

The authors thank the CU Boulder Department of Biochemistry Shared Instruments Pool core facility (RRID:SCR_018986), Annette Erbse, and its staff; Jennifer Doudna and Ben Adler for generously sharing CRISPR-Cas13 editing plasmids; Connor Keane for assistance with Python; Breck Duerkop’s lab and Shelby Anderson for sharing of lab space to complete sample preparation for EM imaging at CU Anschutz; the staff of the cryo-EM facility at UC San Diego, including M. Matyszewski and I. Kuschnerus for facilities and scientific and technical assistance; Marcelo Sousa for helpful discussions and insights; and members of the Whiteley lab for their advice and helpful discussion. This work used the Alpine high performance computing resource at the University of Colorado Boulder.Alpine is jointly funded by the University of Colorado Boulder, the University of Colorado Anschutz, and Colorado State University and with support from NSF grants OAC-2201538 and OAC-2322260. Cryo-electron microscopy data were collected at the UCSD cryoEM Facility, which was built and equipped with funds from UCSD and an initial gift from the Agouron Institute. This study was supported in part by the National Institutes of Health P30CA06934 funded Mass Spectrometry Proteomics Shared Resource SCR_021988. This study was supported in part by the Electron Microscopy Core Facility at CU Anschutz. Data were collected using the JEOL JEM-120i, 120 kV TEM, supported by NIH grant 1S10OD036258-01. Light microscopy was performed at the BioFrontiers Institute’s Advanced Light Microscopy Core (RRID: SCR_018302).

## Author contributions

Conceptualization: LKR, EMK, and ATW. Methodology: LKR, EMK, AD, EGA and ATW. Software: LKR, EMK, and LF. Investigation: all authors. Writing – original draft: LKR, EMK and ATW. Writing – review & editing: all authors. Visualization: LKR, AD and EMK. Supervision: KDC, JP, and ATW. Funding acquisition: KDC, JP, and ATW.

## Data and materials availability

All data are available in the main text or the supplementary materials. Strains, phages, and plasmids used in this study are available upon request.

## Supplementary Materials

Materials and Methods

PDB EM Validation Report

Figures S1–S7

Tables S1–S5

## Experimental Model and Subject Details

### Bacterial Strains and culture conditions

All *E. coli* strains used in this study are listed in **Table S3**. *E. coli* were grown as described previously (*15*). Briefly, bacteria were cultured in LB medium (1% tryptone, 0.5% yeast extract, and 0.5% NaCl) shaking at 37 °C and 220 rpm in 1-3 mL of media in 14 mL culture tubes, unless otherwise indicated. When applicable, chloramphenicol (20 µg/mL) or carbenicillin (100 µg/mL) were added. Constructed strains were frozen for storage in LB with 30% glycerol at - 70 °C. *E. coli* OmniPir was used for construction and propagation of all plasmids (*15*). *E. coli* (CGSC6300) was used to collect all experimental data. *E. coli* BL21 (DE3) (NEB cat#: C2527H) or *E. coli* Rosetta2 with the pRARE2 plasmid (Millipore Sigma cat#: 71400-3) were used to express proteins for purification.

When indicated, experimental data was collected using MMCG medium (47.8 mM Na_2_HPO_4_, 22 mM KH_2_PO_4_, 18.7 mM NH_4_Cl, 8.6 mM NaCl, 22.2 mM Glucose, 2 mM MgSO_4_, 100 µM CaCl_2_, 3 µM Thiamine, Trace Metals at 0.1× ((Trace Metals Mixture T1001, Teknova; final concentration: 8.3 μM of FeCl_3_, 2.7 μM of CaCl_2_, 1.4 μM of MnCl_2_, 1.8 μM of ZnSO_4_, 370 nM of CoCl_2_, 250 nM of CuCl_2_, 350 nM of NiCl_2_, 240 nM of Na_2_MoO_4_, 200 nM of Na_2_SeO_4_ and 200 nM of H_3_BO_3_)). When experiments using MMCG required bacteria expressing two plasmids, strains were grown using reduced antibiotic concentrations (MMCG with 20 µg/mL carbenicillin and 4 µg/mL chloramphenicol).

### Phage Amplification and Storage

All phages used this study are listed in **Table S4**. Phages were amplified via either liquid or plate amplification following the protocol for a modified double agar overlay (*50*). For liquid amplification, 5 mL mid-log cultures of *E. coli* in LB plus 10 mM MgCl_2_, 10 mM CaCl_2_, and 100 µM MnCl_2_ were infected with phage at an MOI of 0.1 and grown, shaking, for 2–16 hours. Cultures were then pelleted to remove cellular debris. The supernatant was harvested by decanting and ∼100 µL chloroform was added to the cultures to remove any bacterial contamination.

For plate amplification, 400 µL of mid-log were mixed with 3.5 mL LB soft agar mix (LB with 0.35% agar and 10 mM MgCl_2_, 10 mM CaCl_2_ and 100 µM MnCl_2_) and 100-1,000 PFU. Plates were then incubated for 16 hours at 37 °C. 5 mL of SM buffer (100 mM NaCl, 8 mM MgSO_4_, 50 mM Tris-HCl pH 7.5, 0.01% gelatin) was added to the plate and allowed to soak out the phages for 1 hour before SM buffer was collected and passed through a 0.2 µm filter or treated with 1–3 drops of chloroform to remove viable bacteria. All phages were stored at 4 °C in SM buffer (100 mM NaCl, 8 mM MgSO_4_, 50 mM Tris-HCl pH 7.5, 0.01% gelatin) or LB.

## Method Details

### Generation of Representative Phage gene clusters

Representative phage gene clusters were generated using all Tequatrovirus family reference genomes from NCBI, 16 *Tequatrovirus* genome sequences found in the BASEL collection (*51*), and the genomes representing the common lab phages T2, T4, and T6, for a total of 91 phage genome sequences. A complete list of the input genomes used can be found in **Table S1**. The CDS regions for each of these genomes were extracted using Geneious. Excluding proteins that were less than 50 amino acids, we used the MMSeq2 clustering algorithm (*21, 22*) (default parameters: 80% identity and 80% coverage) to cluster the remaining 22,719 proteins into 862 representative clusters. Any clusters representing only one input protein were removed, leaving us with a final set of 629 phage proteins representing those commonly expressed by phages in the genus *Tequatrovirus*. For a full list of the proteins represented in each cluster, see **Table S1**.

### MSA generation and protein structure prediction

Multiple sequence alignments for each gene were generated using AlphaPulldown (*52*). Generated MSAs were then used as input for AlphaFold-multimer (*53*). In AlphaFold, each model generated three predictions for a total of 15 predicted structures per interaction. After the predictions were completed, the ipTM + pTM scores (“weighted pTM”) for each model were averaged, and the average weighted pTM score was used to identify confidently predicted interactions for further follow-up. The top eight hits with homologs in either phage T2 or T4 were selected for additional experimental validation.

### Plasmid construction

The plasmids used in this study are listed in **Table S3**. DNA manipulations and cloning were performed as previously described (*15, 54*). Briefly, target genes were amplified from plasmid, phage or bacterial genomic DNA using Q5 Hot Start High Fidelity Master Mix (NEB, M0494L) flanked by 18 base pairs of homology to the vector backbone. Vectors were digested using restriction digest and genes were ligated into vectors using modified Gibson Assembly (*55*). Gibson reactions were then transformed via heat shock or electroporation into competent OmniPir (*15*) and plated onto appropriate antibiotic selection. When possible, phage gene coding sequences were amplified from the genomic DNA of the indicated *E. coli* phages. bNACHT and phage protein point mutations were generated by amplifying the gene of interest in two parts from a plasmid template, with the desired mutation occurring in the overlapping region between the two amplicons. Unless otherwise indicated, all enzymes were purchased from New England Biolabs.

During the process of data collection, the pLOCO2 backbone was modified to reduce extraneous regions, and the resulting backbone was named pLOCO3.

Figures 1D, 4J, S3N, use vectors generated from pLOCO2. Figures 1B, 1E, 3D, 6A, 6E, 6F, 6G, S2B, S5D, S7A, S7B, S7C, S7D, S7D, S7E, S7F use vectors generated from pLOCO3.

For all vectors using the pLOCO2 backbone, pEK0252 was amplified and purified from OmniPir. Purified plasmid was then linearized using SbfI-HF and NotI-HF.

For all vectors using the pLOCO3 backbone, pHL0151 was amplified and purified from OmniPir. Purified plasmid was then linearized using SbfI-HF, NotI-HF, and Sphl-HF.

For all vectors using the pTACxc backbone, pAW1608 was amplified and purified from OmniPir. Purified plasmid was then linearized using BmtI-HF and NotI-HF.

For all vectors using the pETSUMO2 backbone, pAW1642 was amplified and purified from OmniPir. Purified plasmid was then linearized using BamHI-HF and NotI-HF.

For all vectors using the pET backbone, pAW1642 was amplified and purified from OmniPir. Purified plasmid was then linearized using NdeI-HF and NotI-HF.

For all vectors using the pBAD30 backbone, pAW1640 was amplified and purified from OmniPir. Purified plasmid was then linearized using EcoRI-HF and NotI-HF.

Plasmid pEK0278 was purchased from TWIST Biosciences and cloned into the BamHI and XhoI restriction sites of the pET21(+) vector. When needed, DNA sequences were randomly generated to ensure that inserts had a minimum length of 300bp, per manufacturer suggestions.

For constructing vectors encoding the eLbuCas13a system (*56*) with appropriate spacers, the backbone was amplified from pBA681 using PCR (forward primer oEK0228: ATGCTTGGGCCCGAA. Reverse primer oEK0229: GGGCGGAGCCTATGGAAAAACGGCTTTGCCGCG). Gibson ligation was used to circularize the vector with the new insert.

Sanger sequencing (Azenta, Quintara) was used to validate the correct sequence within the multiple cloning site. When needed, nanopore sequencing (Quintara, Plasmidsaurus) was used to obtain the sequence of the entire plasmid.

### Efficiency of plating/phage replication analysis

Efficiency of plating (EOP) was used to determine phage titer and replication. To do this, we employed a modified double agar overlay assay (*50*). Briefly, overnight cultures of *E. coli* expressing the indicated plasmids in MMCG plus appropriate antibiotics were diluted 1:10 into the same media and cultivated for an additional two to three hours to reach logarithmic growth phase (OD600 0.1– 0.8). 400 µL of the culture was then mixed with 3.5 mL 0.35% agar MMCG, plus an additional 5 mM MgCl_2_ and 100 mM MnCl_2_. The mixture was then poured onto a 1.6% agar MMCG plate and cooled for 15 minutes. 2 µL of a phage dilution series in SM buffer was spotted onto the overlay and allowed to adsorb for 10 minutes before the plate was incubated overnight at 37 °C.

Plaque formation was analyzed the following day. Instances with a hazy zone of clearance rather than individual plaque formation at the lowest phage concentration was enumerated as ten plaques. 0.9 plaques at the least dilute spot were used as the limit of detection in instances where no zone of clearance or plaque formation was visible.

### Transformation efficiency analysis

The impact of coexpression of bNACHT alleles and phage proteins on bacterial growth was quantified using a cotransformation assay. Briefly, electrocompetent *E. coli* expressing either bNACHT11 or an empty vector (GFP) was transformed with 50 ng of each purified plasmid encoding the indicated phage protein and recovered for 1 hour in 1 mL SOC (2% Tryptone, 0.5% yeast extract, 10 mM NaCl, 2.5 mM KCl, 10 mM MgCl_2_, 10 mM MgSO_4_, 20 mM glucose). Each transformation was diluted 1:10 and 80 µl of each dilution was spread onto LB plates plus selective antibiotics and 500 µM IPTG. Plates were incubated overnight at 37ºC and quantified the next morning.

### Colony formation/growth inhibition and analysis

The impact of coexpression of different alleles of bNACHT and phage proteins on bacterial growth was quantified using a colony formation assay. Briefly, *E. coli* was cultivated overnight in LB with appropriate antibiotics. Cultures were diluted in a 10-fold series into LB and 5 µL of each dilution was spotted onto an LB agar plate containing the appropriate antibiotics, as well as IPTG as indicated. For osmotic sensitivity assays, dilutions were spotted onto plates containing LB + the specified concentration of NaCl, KCl, or sucrose, appropriate antibiotics, and IPTG as indicated. Spotted bacteria were allowed to dry for 10 minutes before the plates were incubated overnight at 37 °C.

Growth inhibition was measured the following day by enumerating the colony forming units of each strain, reported as CFU/mL for the starting culture. For instances where bacteria were growing but no individual colonies could be counted, the lowest bacterial concentration at which growth was observed was counted as ten CFU. In instances where no growth was visible, 0.9 CFU at the least dilute spot was used as the limit of detection.

### Construction of phage gene deletions

Phage T4 knockout mutations were generated following the protocol described in Adler et al., 2022 (*56*). Briefly, wild-type phages were amplified via either plate or liquid culture (see above) on *E. coli* strains expressing a pET vector encoding the template for homologous repair. Phage lysates amplified in this way were then mixed with 400 µL mid-log bacteria expressing eLbuCas13a constructs with spacers targeting the gene of interest and poured onto an MMCG agar plate as described in solid plate amplification detailed above. Individual plaques were plaque-purified by using a glass Pasteur pipet to extract the plaque from the soft agar, and spot-plated onto *E. coli* expressing the same spacer to confirm that phages were able to evade targeting by CRISPRCas13. Deletion of the target gene was validated by PCR. Validated phage T4 mutants were subsequently amplified via either liquid or plate amplification on *E. coli* in MMCG.

The homologous repair templates were designed to encode 250 bp on either side of the target gene, and the first and last 6 amino acids of the target gene were maintained in the knockout to minimize polar effects. Two 31-nt spacers were selected to target the beginning of each gene.

### Validation of protein expression by western blot

To analyze the expression of bNACHT and phage protein alleles, 3 mL of *E. coli* expressing the indicated plasmid were grown to mid-logarithmic phase in MMCG and 5 × 10^8^ CFU were pelleted. Bacterial pellets were resuspended in 100 µL of 1 × LDS buffer (106 mM Tris-HCl pH7.4, 141 mM Tris Base, 2% w/v Lithium dodecyl sulfate, 10% v/v Glycerol, 0.51 mM EDTA, 0.05% Orange G). Samples were incubated at 95 °C for 10 minutes followed by a 5-minute centrifugation at 20,000 × g to remove debris. Samples in LDS were loaded at equal volumes and resolved using SDS-PAGE, then transferred to PVDF membranes charged in methanol. Membranes were blocked in Licor Intercept Buffer for one hour at 25 °C, followed by incubation with primary antibodies diluted in Intercept buffer overnight at 4 °C with rocking.

α6×His antibody (Thermo) was used at 1:5,000 to detect bNACHT11-6×His and α*E. coli* RNA polymerase B antibody (Biolegend) was used at 1:5,000 as a loading control. For analysis of Co-IP samples, αFLAG antibody (Sigma) was used at 1:15,000 and αVSV-G antibody (Rockland) was used at 1:10,000. Blots were then incubated with Licor infrared (800CW/680RD) αRabbit/Mouse secondary antibodies at 1:40,000 dilution in TBS-T (0.1% Triton-X) for two hours at room temperature and visualized using a Licor Odyssey CLx. Representative images were assembled using Adobe Illustrator CC 2024.

### Protein Alignments

N-terminal bNACHT11 alignments used for the sequence logo in Figure 6C were generated used the NCBI protein-protein BLAST plugin on Geneious to collect the top 500 BLAST results for representative clade 14 bNACHT proteins (*15*): bNACHT01, bNACHT02, bNACHT11, bNACHT12, and bNACHT23, which generated a list of 600 nonredundant protein sequences (*57*). We extracted the first 8 resides from each sequence and aligned them using the Clustal Omega Geneious plugin with default settings. The alignment was manually inspected and sequences with incomplete N-termini or non-identifiable start codons were excluded. The final alignment used for the sequence logo included 537 protein sequences. Sequence logos were generated using WebLogo (https://weblogo.berkeley.edu/logo.cgi). A custom color scheme was used which grouped the following amino acids together: [AILMV], [DE], [NSTQ], [P], [G], [FWY], [HKR].

SNaCT alignments used for coloring the SNaCT domain by conservation in Figure 3K were generated using the NCBI protein-protein BLAST plugin on Geneious to search for bNACHT11 homologs of the SNaCT domain. This BLAST yielded 96 sequences which were aligned using the Clustal Omega Geneious plugin with default settings. This alignment was uploaded to Consurf which was used to color the structure by conservation (*58*).

Alignments used for Figure S3 were generated by selecting the specified sequences and generating an alignment using a MUSCLE 5.1 alignment with default parameters in Geneious. For visualization, Jalview was used to color residues by percent identity.

### Protein expression

Vectors for expressing gp6-6×His and gp6^I20A^-6×His were transformed into BL21 (DE3) and plated onto 1.6% LB agar plates with 100 µg/mL carbenicillin. An individual colony was picked the following day and inoculated into 5 mL of liquid LB media plus 100 µg/mL carbenicillin. The culture was then grown overnight shaking at 37 °C and 220 rpm. The following day, the culture was used to inoculate 0.5–1 L of the same media, then grown to an OD600 of ∼0.6 before IPTG was added to 500 µM to induce protein expression. The culture was then moved to a 16 °C shaking incubator and allowed to grow overnight.

The vector for expressing 6×His-SUMO-bNACHT11 was transformed into Rosetta2 expressing the pRARE2 plasmid and plated onto 1.6% LB agar plates + 100 µg/mL carbenicillin and 20 µg/mL chloramphenicol. An individual colony was picked the following day and inoculated into 5 mL of liquid LB media plus 100 µg/mL carbenicillin and 20 µg/mL chloramphenicol. The culture was then grown overnight shaking at 37 °C and 220 rpm. The following day, the culture was used to inoculate 0.5–1 L of the same media, then grown to an OD_600_ of ∼0.6 before IPTG was added to 500 µM to induce protein expression. The culture was then moved to a 16 °C shaking incubator and allowed to grow overnight.

Vectors for expressing gp9-6×His, gp9I^34A^-6×His, gp57-6×His, and 6×His-SUMO-bNACHT11^I553A^ were transformed into Rosetta2 expressing the pRARE2 plasmid and plated onto 1.6% MMCG agar plates + 100 µg/mL carbenicillin and 20 µg/mL chloramphenicol. An individual colony was picked the following day and inoculated into 20 mL of M9ZB media (47.8 mM Na_2_HPO_4_, 22 mM KH_2_PO_4_, 18.7 mM NH_4_Cl, 85.6 mM NaCl, 1% Casamino acids (VWR), 0.5% v/v Glycerol, 2 mM MgSO_4_, Trace Metals at 0.5 × (Trace Metals Mixture T1001, see above) plus 100 µg/mL carbenicillin and 20 µg/mL chloramphenicol. The culture was then grown overnight shaking at 37 °C and 220 rpm. The following day, the culture was used to inoculate 0.5–1 L of the same media to an OD_600_ of 0.05, then grown to an OD600 of ∼1.5. Cultures were crash-cooled on ice for 20 minutes before IPTG was added to 500 µM to induce protein expression. The culture was then moved to a 16 °C shaking incubator and allowed to grow overnight. Proteins expressed in this way were purified and used in microscale thermophoresis experiments.

### Protein purification

After overnight induction with IPTG, cultures were harvested by centrifugation for 30 minutes at 5,000 rpm and 4 °C in an Avanti JXN-26 Floor Centrifuge using the JXN 12.500 rotor (Beckman). The resulting pellets were resuspended in 40 mL Lysis buffer (20 mM HEPES pH 7.5, 400 mM NaCl, 10% v/v Glycerol, 30 mM Imidazole, 0.1 mM Dithiothreitol (DTT)). After resuspension, cells were lysed by sonication at 80% amplitude, with 30 second pulses for a total processing time of 10 minutes using a Sonicator 4000 (Misonix). Debris was removed from sonicated lysates by centrifugation for 60 minutes at 4 °C and 14,000 × g in a 5910 R centrifuge (Eppendorf). The soluble lysate was then decanted and protein was purified using immobilized metal affinity chromatography. Briefly, the soluble lysate was run over 1 mL of Ni-NTa resin (Fisher Sci) equilibrated in Lysis Buffer. The resin was then washed with 2 × 25 mL of Wash Buffer (20 mM HEPES pH 7.5, 1 M NaCl, 10% v/v glycerol, 30 mM Imidazole, 0.1 mM DTT) and protein was eluted in 5-10 mL of Elution Buffer (20 mM HEPES pH 7.5, 400 mM NaCl, 10% v/v glycerol, 300 mM Imidazole, 0.1 mM DTT). Proteins were then dialyzed against 2 × 1 L of Dialysis Buffer (20 mM HEPES pH 7.5, 250 mM KCl, 0.1 mM DTT), overnight at 4 °C using either 10 kDa MWCO tubing (VWR), 3 kDa MWCO Snakeskin Dialysis Tubing (VWR), or 3 kDa MWCO Slide-A-Lyzer Dialysis cassettes (VWR) as appropriate for the molecular weight of the protein.

The 6×His-SUMO-tag was cleaved from alleles of bNACHT11 using 6×His-hSENP2 (produced in-house) during the overnight dialysis step. After dialysis, proteins were run over 1 mL Ni-NTA beads equilibrated in dialysis buffer to remove any uncleaved 6×His-SUMO tagged proteins.

After dialysis, proteins were concentrated as needed using 3 kDa or 30 kDa MWCO Nanosep spin concentration columns (Pall Labs) and stored 200–500 µL aliquots in Dialysis Buffer at –70 °C. Protein concentrations were measured using A_280_ on a Nanodrop OneC (Thermo) and protein purity was visualized using SDS-PAGE followed by Coomassie staining. Proteins purified in this way were used in microscale thermophoresis experiments.

### Microscale thermophoresis

Protein binding affinities were measured using microscale thermophoresis (*59*). For these experiments, alleles of phage proteins containing C-terminal 6×His tags were labeled using a Monolith His-Tag Labeling Kit RED-tris-NTA 2nd Generation (NanoTemper: Cat# MO-L018) following manufacturer instructions. Samples were allowed to equilibrate for 30 minutes at room temperature before MST measurement. Experiments were performed using independently labeled proteins and independently pipetted ligand titrations. Measurements were performed using 60% laser excitation, medium MST power, and a chamber temperature of 25 °C on a Nano-BLUE/RED Monolith NT.115 (NanoTemper). All data was analyzed using a hot time of 9–10 seconds. Fraction bound values were calculated by Mo.AffinityAnalysis software (NanoTemper). Binding data from three independent experiments were fit using the quadratic binding equation (**Figure S3H-L**) (*60*).

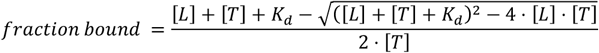

In this equation [L] is the concentration of ligand, [T] is the concentration of labeled target (50 nM for all experiments reported here). The dissociation constants reported here represent the average of the dissociation constants calculated for each biological replicate ± standard error of the mean.

All reactions used MST Buffer (20 mM HEPES pH 7.5, 250 mM KCl, 5 mM MgCl_2_, 0.05% v/v Tween-20, 0.1 mM DTT, 1 mM ATP). For high-salt measurements, salt concentrations were increased to 1 M KCl.

### Oligomerization Analysis

To express 6×His-SUMO-bNACHT11, gp6-6×His, gp9-6×His, and 6×His-SUMO-gp57, the respective constructs were transformed into *E. coli* Rosetta2 pLysS (EMD Millipore). The primary cultures (10 mL) were grown in the presence of appropriate antibiotics (see **Table S3** for construct details and antibiotic usage). The following day, primary cultures were used to inoculate 1 L of 2XYT medium (1.6% tryptone, 1% yeast extract, 0.5% NaCl) in 2 L flasks and grown at 37 °C until the OD_600_ reached approximately 0.75. Protein expression was induced by adding 0.33 mM IPTG and the cultures were incubated overnight at 20 °C to promote protein production. After 14–16 hours, the cells were harvested by centrifugation, and the bacterial pellets were resuspended in ice-cold resuspension buffer (50 mM Tris-HCl (pH 7.5), 300 mM NaCl, 10 mM imidazole, 10% glycerol, and 2 mM β-mercaptoethanol). Resuspended cells were lysed using a sonicator and the lysate was clarified by centrifugation. Proteins were purified through Ni^2+^ affinity chromatography (Ni-NTA Superflow, Qiagen). For bNACHT11 and gp57, the N-terminal 6×His-SUMO tags were removed by buffer exchange to eliminate imidazole using a buffer containing 50 mM Tris (pH 7.5),300 mM NaCl, 10% glycerol, and 2 mM β-mercaptoethanol, followed by digestion with SENP2 overnight at 4 °C. The cleaved tags were removed the next day by passing the proteins through Ni^2+^ beads again. All purified proteins were concentrated and further purified using a Superdex 200 Increase 10/300 GL size exclusion column (Cytiva) in a buffer containing 20 mM HEPES (pH 7.4), 200 mM KCl, and 1 mM dithiothreitol (DTT). The purity of the proteins was confirmed by analyzing the samples using SDS-PAGE, followed by staining with Coomassie blue (**Figure S3B-F**).

To study the effect of phage trigger addition on the oligomeric state of bNACHT11, size exclusion chromatography was performed. Briefly, 25 µM bNACHT11 was incubated with or without 1 mM ATP, 1 mM ATPγS, 1 mM AMP-PNP, or 1 mM ADP in a buffer containing 20 mM HEPES (pH 7.4), 200 mM KCl, 10 mM MgCl_2_, and 1 mM DTT. The reaction mixtures were incubated at room temperature for 2–3 minutes. A 2–3 molar excess of either gp6, gp9, or gp57 purified protein was then added to the reaction mixture, and the samples were further incubated at room temperature for 5 minutes. The samples were immediately transferred to ice and kept on ice until injected into a Superdex 200 Increase 10/300 GL size exclusion column (using buffer 20 mM HEPES pH 7.4, 200 mM KCl, 10 mM MgCl_2_, and 1 mM DTT). The corresponding protein fractions were collected in a 96-well block, changes in the elution volume were analyzed, and indicated samples from the peaks were subjected to SDS-PAGE analysis.

### CryoEM sample preparation

For single-particle analysis, bNACHT11 and gp57 proteins were purified as described in the sections above, “*Protein Expression” and “Protein Purification*”. The samples were prepared immediately before grid preparation. Briefly, the protein samples were mixed in a PCR tube at 25 µM and 75 µM concentrations for bNACHT11 and gp57, respectively. Next, ATP was added to the mixture so that the final sample buffer composition is 20 mM HEPES, 200 mM KCl, 1 mM DTT, 5 mM MgCl_2_, and 2 mM ATP. The sample was incubated for 10 minutes at room temperature and then stored on ice until grid preparation.

Initial extensive efforts, which included changing the buffer composition and inclusion of detergents, were futile as the resulting sample showed strong preferred orientation. To address the preferred orientation issue, we chose to prepare the grids using the SPT Labtech Chameleon instrument, which offers features like blot-free sample application and high-speed plunge freezing. In the Chameleon (SPT Labtech), liquid ethane was maintained at approximately -175 °C, and the humidity was set to be greater than 75%. “Self-wicking” grids (Quantifoil Active 300 mesh) were loaded into the instrument and glow discharged for 30 seconds at 12 mA.

To prepare the grids, 7 µL of sample was aspirated into the dispenser and subsequently tested for proper dispensing within the Chameleon software before being deposited onto the grid. Based on the videos recorded by the instrument, we prepared three grids. The grid used for data collection had a 255 ms sample plunging time, and the sample was vitrified in liquid ethane and stored in liquid nitrogen dewar until loaded onto the electron microscope.

### CryoEM data collection and data processing

CryoEM grids were securely mounted into standard AutoGrids (ThermoFisher Scientific) for imaging. The imaging was done using a Titan Krios G4 transmission electron microscope (ThermoFisher Scientific), operated at a voltage of 300 kV. This microscope was configured for fringe-free illumination and equipped with a Falcon 4 direct electron detector, complemented by a Selectris X energy filter. The microscope was operated in EFTEM mode with a slit-width of 20 eV. Automated data acquisition was performed using EPU (ThermoFisher Scientific). A sample tilt was applied during the data collection, resulting in a roughly similar number of movies across a range of tilted angles from 0° to 35°, with each angle incrementation as 10°, 20°, or 35°. These movies were collected at a magnification of 130,000X with a pixel size of 0.935 Å. A total dose of 51 e^-^/Å^2^ was applied at a rate of 7 eps. During the data collection, a defocus range of -0.5 to -2.2 was used. Following these settings, a total of 8,250 movies were collected. After applying various parameters, such as CTF fitting and excessive motion criteria, 7,622 movies were selected for the final data processing.

The cryoEM datasets were processed using cryoSPARC version 4.7.0 (*61*). First, movies were motion-corrected using patch motion-correction (multi) and CTF estimations were performed using CTF estimation (multi) (*62*). For the initial particle picking on 1,000 micrographs data subset, a blob-picking strategy with a diameter ranging from 120 to 200 Å was used. The selected particles were curated using 2D classification, followed by *ab initio* modeling. This was then followed by heterogeneous and homogeneous refinement with C7 point group symmetry. The resulting map was used to generate templates, and the procedure was repeated with 1,000 micrographs once more. This resulted in a high-resolution map, which was further used to generate templates. The entire curated dataset of 7,622 micrographs was then processed. Multiple rounds of 2D classification-based particle cleaning, *ab initio*, and heterogenous refinement resulted in distinct reconstructions that could be visually distinguished from each other. The predominant ring-like classes were selected. To compensate for the relatively lower proportion of particle-side views, a “Rebalance 2D” job was executed. This resulted in 141,477 particles that were used in the reconstruction of a 2.5 Å map using NU-refinement with C7 symmetry (*63*).

Careful analysis of the reconstructed 2.5 Å map revealed that the density for bound gp57 was poor. We reasoned that the poor density of gp57 could be due to the additive contributions from high flexibility in the SNaCT region of bNACHT11, the small size of gp57, and a relatively low occupancy compared to that of the bNACHT11 protomers in the overall complex. To improve the map resolution in the SNaCT region for confident modeling of the bNACHT11 and gp57 complex, we symmetry expanded the particles contributing to the major ring-like reconstruction. This was followed by local refinement at the sensor region using a dynamic mask. Additionally, to separate the particles bound to gp57 from those unbound at the SNaCT domain region, 3D variability analysis was performed. The volume series were manually inspected, and the particle clusters contributing to relatively high gp57 density were selected for further local refinement. The resulting map was used for docking an AlphaFold3-generated model, as discussed below. The overall data processing workflow is shown in Figure S7.

### bNACHT11-gp57 model building and refinement

An initial model for bNACHT11 was generated using AlphaFold server (*64*). First, the “core region” of bNACHT11 (residues 43 to 464) was manually docked into the 2.5 Å map using UCSF ChimeraX (*65*). Subsequently, it was manually built using COOT, ligands were added, and refined in phenix.refine against unsharpened maps, using secondary structure restraints, minimization global and ADP options were selected.

To construct the bNACHT11 SNaCT-gp57 model, we manually docked the AlphaFold3 model of the bNACHT11-gp57 complex into the composite map (deposited as an additional map with EMDB entry EMD-70775) using UCSF ChimeraX. Subsequently, we trimmed the model in COOT to ensure that only the bNACHT SNaCT domain (residues 466-628) and gp57 remained docked in the map. Next, we refined the model using ISOLDE as a plugin for UCSF ChimeraX (*66*). Finally, we refined the model further in COOT, trimming the highly flexible loops with poor density and applying real-space refinement in phenix.refine with rigid body and ADP options selected.

To construct the complete bNACHT11-gp57 complex model, the refined “core region” and “bNACHT11 SNaCT-gp57” models were docked into the 2.5 Å map using UCSF ChimeraX. These models were then merged in COOT, and the missing linker residues were manually built. This monomeric model of bNACHT11 bound to gp57 was subsequently symmetry expanded, chains were renamed, and a rigid body docking was performed for the resulting 7:7 complex. The AlphaFold3 model of bNACHT11 was used as a guide to build the C-terminal tail residue (with poorly resolved but easily observable density) involved in protomer-protomer interactions. Finally, the resulting model was refined again using phenix.refine with rigid body, ADP, and NCS options selected. The Cryo-EM data collection, refinement, and validation statistics are provided in Table S5.

### Structural Data availability

The bNACHT11-gp57 cryoEM reconstruction has been deposited to the Electron Microscopy Data Bank (EMDB) under accession number EMD-70775 and the corresponding coordinate file has been submitted at the RCSB PDB under accession number 9ORF.

### Immunoprecipitation (IP)

*E. coli* containing an IPTG inducible untagged or VSV-G tagged phage protein was grown overnight in selective antibiotic and diluted the next morning 1:26 in 25 mL of LB + antibiotic until they reached mid-logarithmic phase (OD600 ∼0.5). 500 µM IPTG was added and strains were incubated for an additional 30 minutes. OD600 normalized cultures (2.5 × 10^10^ CFU) were then centrifuged at 4,000 ×g for 10 minutes at 4ºC. The resulting pellet was resuspended in 3 mL lysis buffer (20 mM Trix-HCl pH 7.5, 400 mM NaCl, 2% w/v glycerol, 0.1% v/v Triton X-100, and 0.1 mM DTT). Cells were lysed by sonication at 20% amplitude with three 30 second pulses using a Sonicator 4000 (Misonix). Lysate was then centrifuged at 20,000 *× g* at 4ºC. A portion of each clarified lysate was saved as “Input”. For each sample, 50 µl VSV-G agarose beads (Sigma) were prepared by washing three times in 1000 µl lysis buffer. Samples were then mixed with beads overnight at 4ºC with end-over rotation. The next day, beads were washed three times with 10 mL lysis buffer. IP beads were resuspended in 50 µl LDS. 3 µl of each sample was analyzed on a 4-20% SDS-PAGE. Silver staining was performed using SilverQuest silver staining kit (Thermofisher) according to manufacturer’s instructions.

### Mass spectrometry

Peptide extraction of the gel bands was carried out as previously described (*67*). Briefly, gel bands were destained in 25 mM Ammonium bicarbonate. The de-stain was subsequently removed, and samples were reduced in 5 mM TCEP. The bands were then alkylated with 20mM chloroacetamide (CIAA) for 45 min in the dark at room temperature. CIAA was then removed and washed with 25 mM ammonium bicarbonate by shaking for 10 min at room temperature. The samples were again washed with 25 mM ammonium bicarbonate in 50% acetonitrile by vortexing for 10 min at room temperature. Next, the samples were dehydrated by treating with 100% acetonitrile for 10 min without vortexing. The samples were then dried by vacuum concentration for 10 min at room temperature. Samples were rehydrated with 100 µL of cold (4 °C) trypsin (10 ng/µL) and left at 4 °C for 45 min. The gel pieces were then incubated at 37 °C overnight to complete the digestion. Twenty µL/sample of 5% formic acid (FA) was added to stop digestion. The supernatant was transferred to new tubes. Eighty μL/sample of 50% acetonitrile/0.1%FA was then added, and the samples were vortexed for 20 min at room temperature. The supernatant was again transferred to the collection tubes. Eighty µL/sample of 50% acetonitrile/0.1%FA was added to the collection tubes, and the samples were vortexed for 15 min at room temperature. Samples were dried down via vacuum concentration. The samples were then re-suspended in 0.1% FA for mass spec analysis.

Samples were analyzed on a TIMS Q-TOF mass spectrometer (Bruker Daltonics) coupled with an Evosep One liquid chromatography-mass spectrometry (LC-MS) interface as previously described. Data were analyzed with FragPipe version 22.0, utilizing the following parameters. Precursor tolerance was set as ±15 ppm and fragment tolerance as ±25 ppm. Data was searched against the *Escherichia coli* database UniProt ID UP000000625. The database was adopted with 50% decoys and the MSFragger common contaminants. All searches were carried out with trypsin enzymatic cleavage. Variable modifications were set as methionine oxidation (15.9949) and N-terminal acetylation (42.016). The fixed modification was set as cysteine (57.021465). Match between runs was utilized within MS1 quantification. Searches were performed with all replicates from each sample condition grouped to obtain protein coverage across all analyzed replicates. Results were filtered to 1% FDR at the peptide and protein levels.

### Co-Immunoprecipitation (Co-IP)

Co-IPs were performed for validation of protein-protein interaction by mass spectrometry. Strains containing IPTG-inducible untagged or VSV-G tagged phage protein and arabinose-inducible 3×FLAG tagged kinase were grown overnight in LB plus appropriate antibiotics and 0.2% arabinose. The next morning, strains were diluted 1:26 in LB plus appropriate antibiotics and 0.2% arabinose until they reached mid-logarithmic phage (OD600 ∼0.5). 500µM IPTG was added and strains were incubated for an additional 30 minutes. Lysate was prepared as described above (see Immunoprecipitation (IP)). For each sample, 60 µl FLAG-magnetic beads (Sigma) were prepared by washing three times in 1000 µl lysis buffer. Samples were then mixed with beads overnight at 4ºC with end-over rotation. The next day, beads were washed three times with 10 mL lysis buffer. IP beads were resuspended in 50 µl LDS. 3 µl of each sample was analyzed by SDS–PAGE and western blots as described above.

### Laser Scanning Confocal Microscopy

Strains were grown overnight in LB plus appropriate antibiotics. The next day, cells were pelleted and resuspended in 20 mL fresh MMCG media. Strains were grown to mid-logarithmic phase (OD_600_ ∼0.5) and induced with 500 µM IPTG. Following 30 minutes of incubation the strains were OD normalized to 0.5 and stained with 1 µg ml^−1^ FM4-64 for two minutes. 5ul of each sample was pipetted onto an MMCG imaging pad containing 500 µM IPTG. Samples were imaged using a Nikon AXR confocal microscope system equipped with an NSPARC detector. Images were acquired using a 100x oil objective and 1.45 numerical aperture. Cells were focused using the AXR detector and the final image was acquired using the NSPARC detector. Images have a pixel size of 0.06 µm/pixel with a final image size of 512 × 512 pixels. The images were acquired sequentially where EGFP signal was measured using an excitation of 488 nm and an emission of 524 nm and then FM4-64 signal was measured using an excitation of 488 and an emission of 699 nm. All images were acquired using the same laser power. Each image was further analyzed using the Richardson-Lucy 2D deconvolution method with 20 iterations.

### TEM Imaging

Strains were grown overnight in LB plus appropriate antibiotics and diluted 1:100 in fresh medium with no antibiotics the next day. Strains were grown until they reached an approximate OD of 0.5 before induction with 500 µM of IPTG. Strains were grown with inducer for 30 minutes before being pelleted and resuspended in fixative solution. Cells were fixed with 2% paraformaldehyde and 2% glutaraldehyde in 0.1 M sodium cacodylate buffer and embedded in 3% agarose. Samples were trimmed into small blocks and rinsed three times in 0.1 M sodium cacodylate buffer, then post-fixed in 2% osmium tetroxide with 0.8% K_3_ [Fe (CN)_6_] in 0.1 M sodium cacodylate buffer for 1 hour at room temperature. Cells were rinsed with water and en bloc stained with 1% aqueous uranyl acetate for 1 hour. Samples were dehydrated through a graded ethanol series, infiltrated with SPURR resin, and polymerized overnight at 70°C. Blocks were sectioned using a diamond knife (Diatome) on a UC7 ultramicrotome (Leica) and collected onto copper grids. Images were acquired on a JEM 120i transmission electron microscope (JEOL) equipped with a LaB_6_ source at 120 kV using a NanoSprint15Mk-II (AMT) sensor.

### Quantification and Statistical Analyses

Data was plotted using GraphPad Prism 10.3.1 at an n of 3 with error bars indicating standard error of the mean. Figures were created using Adobe Illustrator CC 2025 v30.0.

#### NCBI Accession numbers

T2 phage genome: NC_054931.1

T4 phage genome: AF158101.1

T4 CDS 30.5 (gp30.5, T4 p208): NP_049819.1

T2 gp6: YP_010073654.1

T2 gp9: YP_010073657.1

T2 gp21: YP_010073669.1

T2 gp28: YP_010073676.1

T2 gp46: YP_010073694.1

T2 gp57: YP_010073705.1

T2 gp72: YP_010073720.1

bNACHT11 *K. pneumoniae*: WP_114260439.1

bNACHT11 *E. coli*: WP_220676828

HELLP: (CDP29920.1)

Avast2 (WP_063118745.1)

bNACHT01: WP_015632533.1

bNACHT02: WP_021557529.1

bNACHT12: WP_021519735.1

bNACHT23: WP_000433597.1

#### UniProt Identifiers

coaA: P0A6I3

ppK: P0A7B1

glpK: P0A6F3

Additional genome and protein accession numbers for generating non-redundant database of phage proteins to screen in AlphaFold can be found in Table S1.

**Fig. S1.**
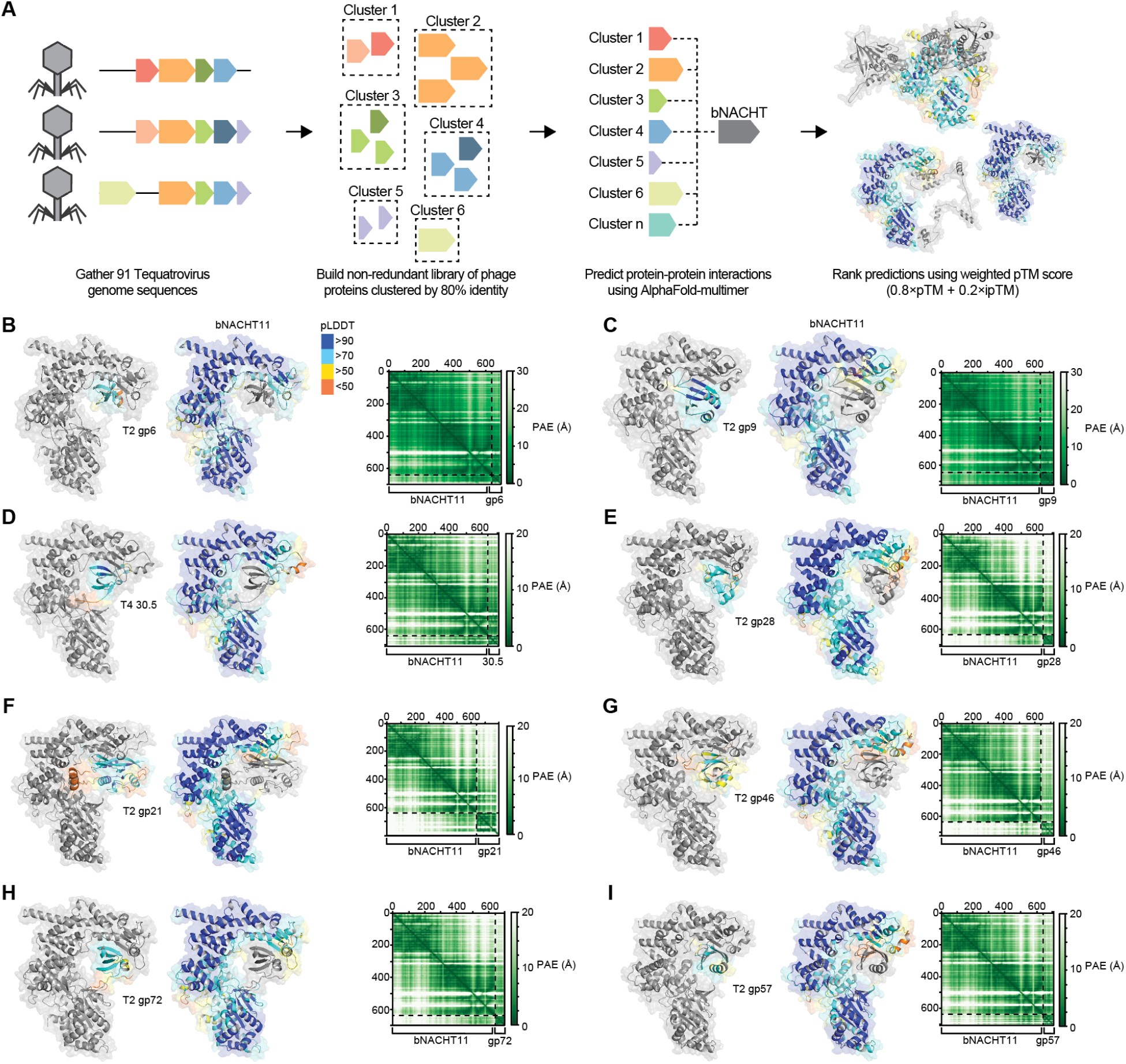
Generation of predicted bNACHT11 activators of bNACHT11, related to Figure 1. **(A)** Schematic of the workflow for an *in-silico* screen to identify phage-encoded activators of clade 14 bNACHT proteins (see Methods). Proteins encoded by 91 phages in the *Tequatrovirus* family were clustered using MMSeq2 (*21, 22*) and 629 representative proteins were then screened using AlphaFold-multimer to predict interactions with bNACHT proteins of interest. See **Table S1** for a list of all proteins used to generate the clusters, and a final list of the cluster representatives. **(B-I)** Predicted structure of bNACHT11 binding the indicated phage proteins. Left: The phage proteins screened are colored according to pLDDT. Middle: bNACHT11 is colored according to pLDDT. Right: Predicted aligned error (PAE) of the predicted interaction between bNACHT11 and the indicated phage protein. (B) T2 gp6. (C) T2 gp9. (D) T4 30.5. (E) T2 gp28. (F) T2 gp21. (G) T2 gp46. (H) T2 gp72. (I) T2 gp57.

**Fig. S2.**
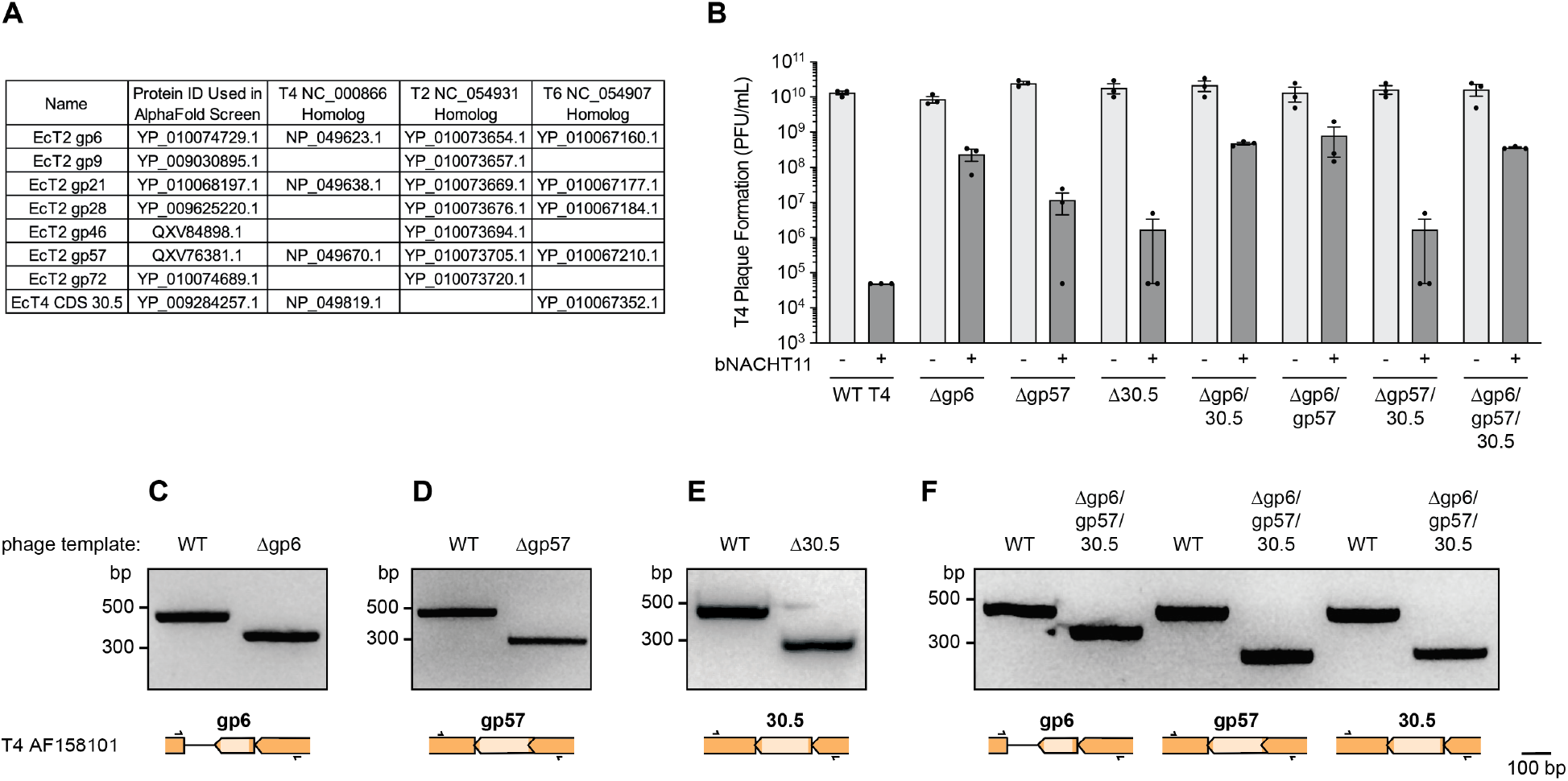
Activator knockouts in phage T4 partially evade bNACHT11 protection. **(A)** Table of homologs of predicted activators among phage T4, phage T2, and phage T6. **(B)** Efficiency of plating of phage T4 of the indicated genotype infecting *E. coli* expressing an empty vector (-) or bNACHT11 (+). Data represent the mean ± SEM of n = 3 biological replicates, shown as individual points. **(C-F)** Validation of knockout generation by PCR and agarose gel separation. (C) Amplification of base pairs 5,552 – 5,986 encompassing gp6 using WT T4 or Δgp6 as a template. (D) Amplification of 34,844 – 35,272 encompassing gp57 using WT T4 or Δgp57 as a template. (E) Amplification of 128,824 – 129,259 encompassing CDS 30.5 using WT T4 or Δ30.5 as a template. (F) Amplification of the specified region using WT T4 or Δgp6/gp57/30.5 as a template.

**Fig. S3.**
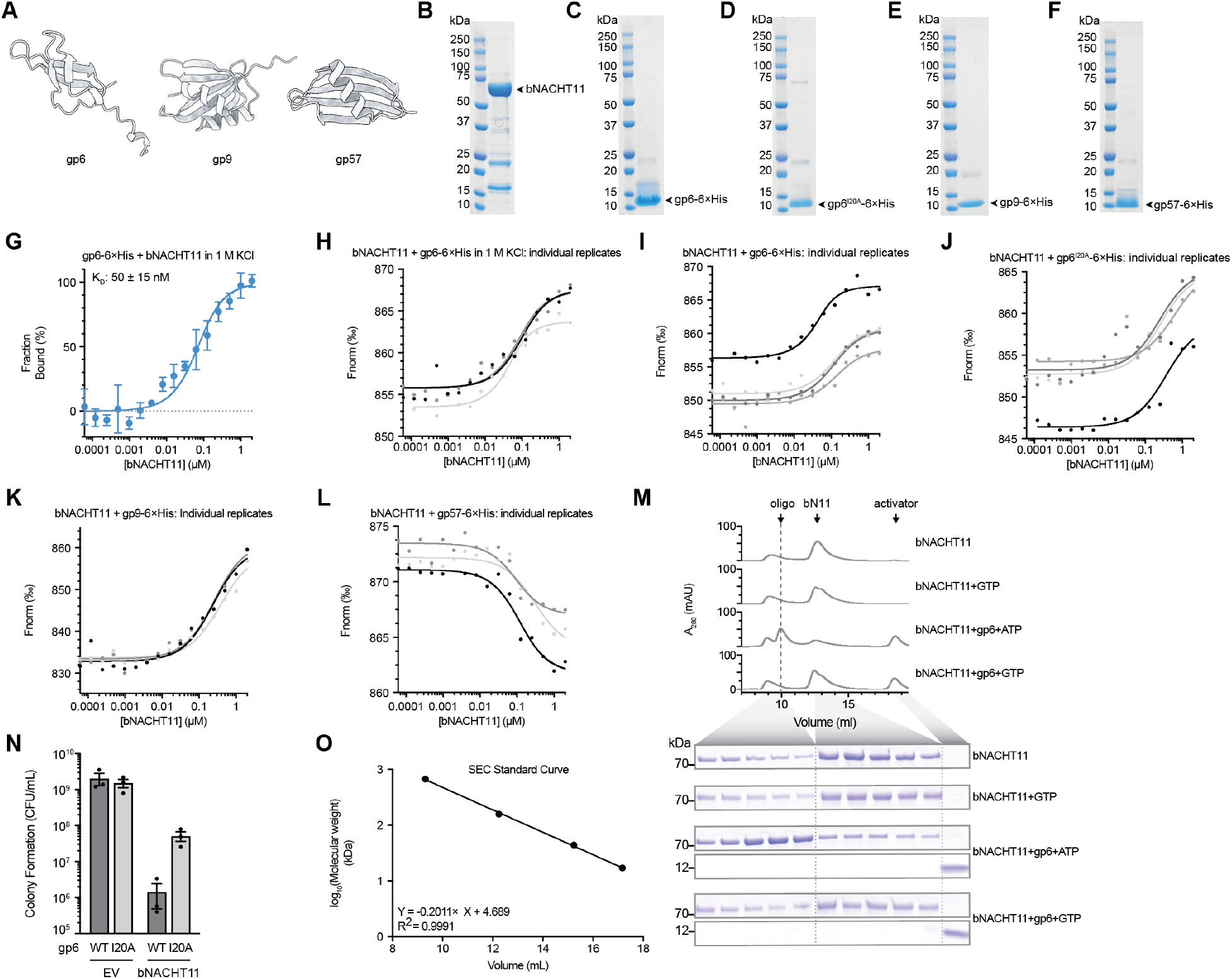
Binding and oligomerization of bNACHT11 with activators, related to Figure 2. **(A)** AlphaFold3 predicted structures of the indicated phage proteins. gp6 pTM = 0.63, gp9 pTM = 0.82, gp57 pTM = 0.80. **(B–F)** Coomassie stained SDS-PAGE gels of the purified protein indicated with an arrow. **(G)** Binding curve of purified His-tagged gp6 and purified bNACHT11, measured using microscale thermophoresis (MST) in 1 M KCl. Data represent the average of n = 3 individual replicates ± SEM. **(H–L)** Normalized fluorescence data for individual MST experiments used to generate the binding curves shown in **Figure S3G** (H), **Figure 2A** (I, J) **Figure 2B** (K) and **Figure 2C** (L). **(M)** Above: Traces showing the absorbance at 280 nM (A_280_) of the indicated samples run over a size exclusion column. Labels indicate the identity of proteins in each peak. Below: Coomassie staining of representative samples taken from the fractions in between the indicated elution volumes. **(N)** Quantification of colony formation of *E. coli* expressing wild-type gp6 (WT) or gp6^I20A^ (I20A) with a C-terminal VSV-G tag on one plasmid and empty vector (EV) or bNACHT11 on a second plasmid. Data represent the mean ± SEM of n = 3 biological replicates, shown as individual points. **(O)** Standard curve of proteins with known molecular weights run over the Superdex 200 Increase size exclusion column used in (M).

**Fig. S4.**
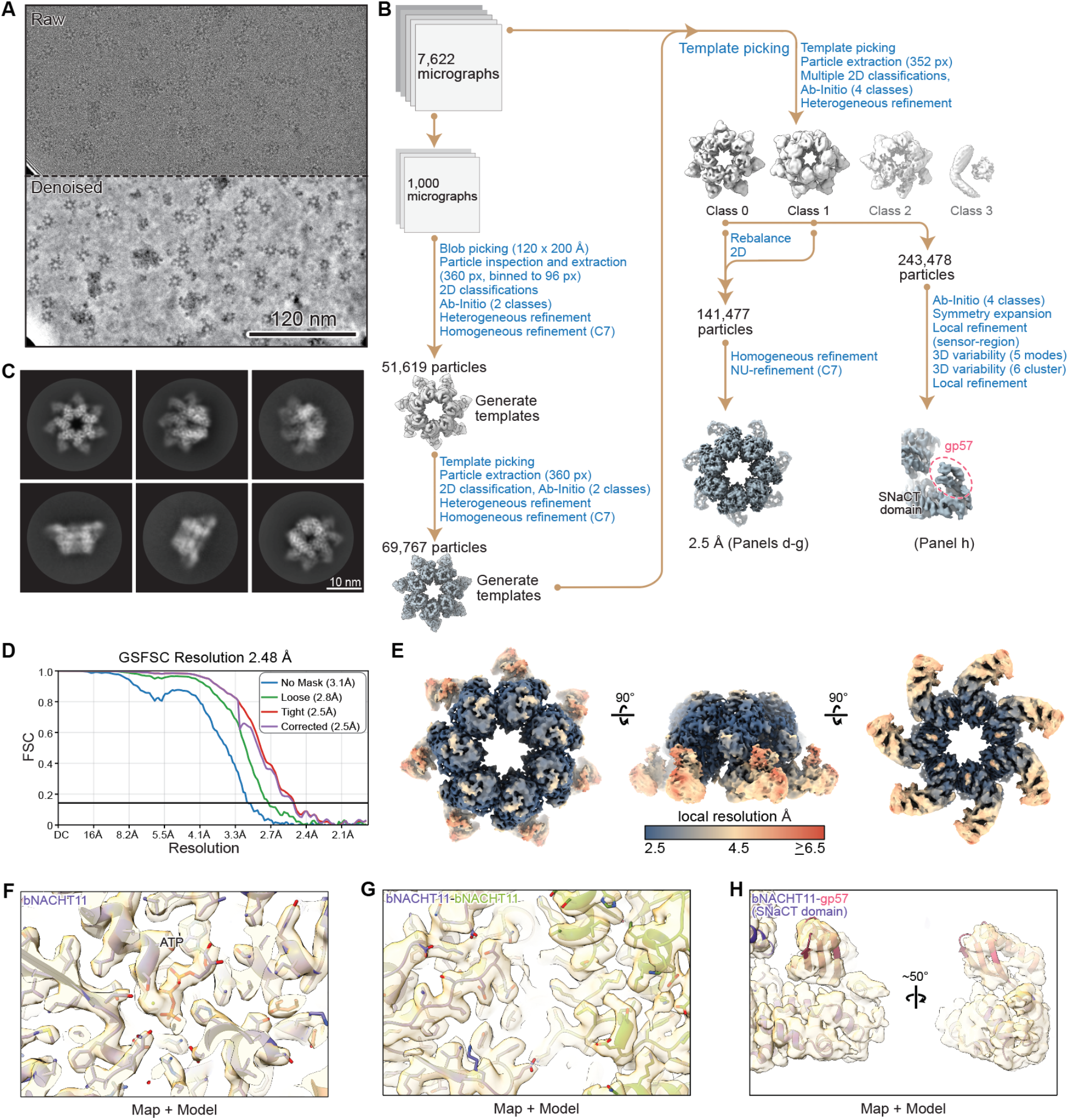
CryoEM workflow for bNACHT11-gp57, related to Figure 4. **(A)** Representative micrograph (from 7,622 micrograph dataset) displaying the bNACHT11-gp57 sample. **(B)** Workflow outlining the cryo-EM reconstruction process for bNACHT11-gp57 using cryoSPARC. **(C)** Representative 2D classes displaying bNACHT11-gp57. **(D)** Gold-standard FSC curve for the final global refinement of bNACHT11-gp57. **(E)** Three views of the globally refined cryo-EM map for the bNACHT11-gp57 heptamer, color-coded by local resolution from 2.5 Å (blue) to >6.5 Å (red). **(F-H)** Example cryo-EM density with a built atomic model demonstrating the model fit. Purple represents one bNACHT11 protomer (F). Green represents an adjacent bNACHT11 protomer (G). Red represents gp57 (H).

**Fig. S5.**
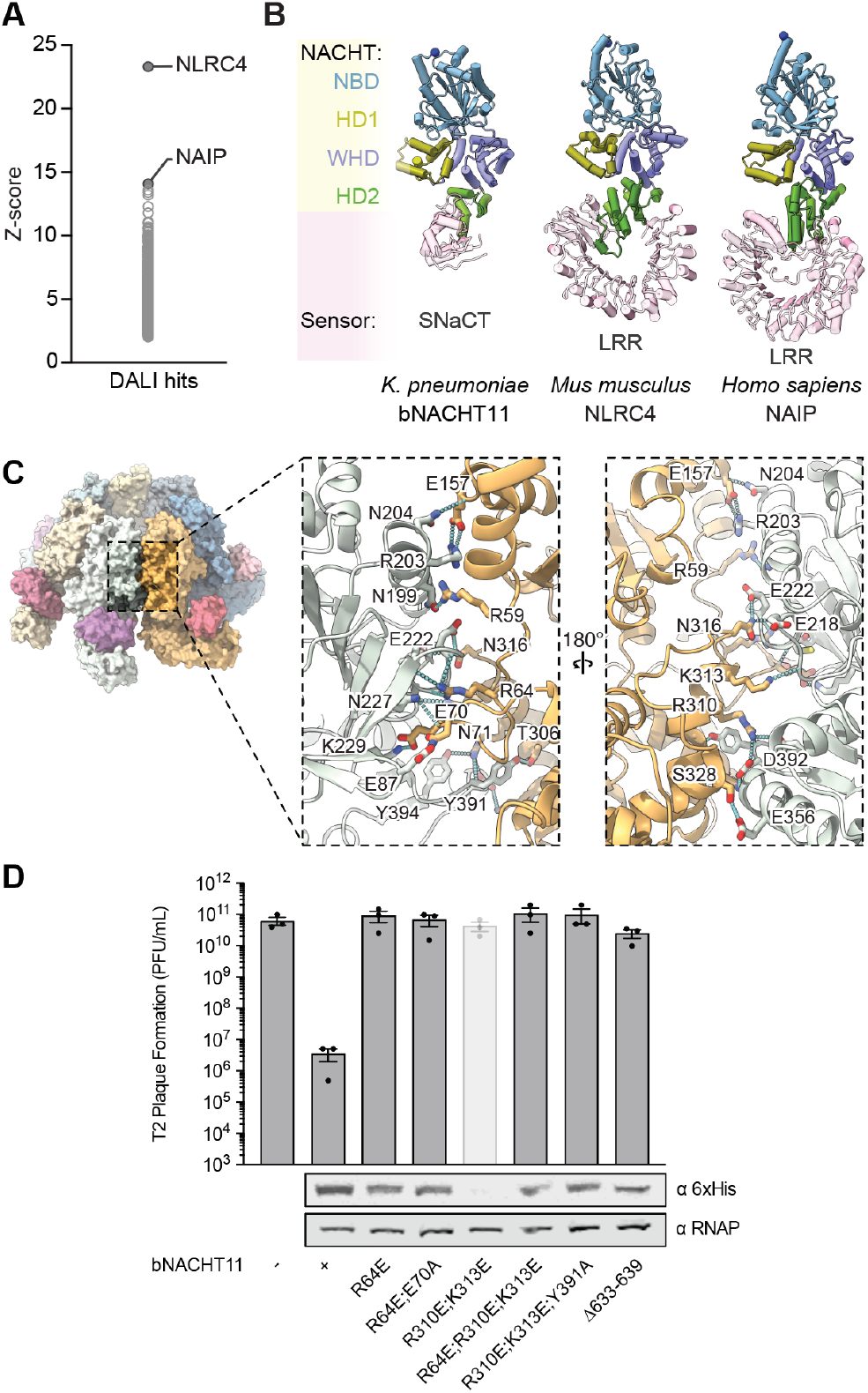
NACHT domain oligomerization disrupting mutations, related to Figure 4. **(A)** Top hits from DALI analysis of bNACHT11 (PDB: 9ORF). **(B)** Top DALI hits for bNACHT11 (PDB: 9ORF): NLRC4 (PDB: 3JBL) and NAIP (PDB: 8FVU) highlighting the conserved NACHT domain and variable sensing domains. **(C)** Assembly interface analysis of gp57-bound bNACHT11 complex. Zoomed in view of NACHT domain assembly interface from bNACHT11-gp57 cryoEM structure. Grey and orange distinguish two bNACHT11 protomers. The residues involved in polar interactions are shown as side-chain sticks representation and are labelled. **(D)** Top: Efficiency of plating of phage T2 infecting *E. coli* expressing an empty vector (-) or the indicated genotype of bNACHT11-6×His. Data represent the mean ± SEM of n = 3 biological replicates, shown as individual points. Mutations resulting in no protein expression are indicated as bars with decreased opacity. Bottom: Western blot analysis of *E. coli* expressing 6×His-tagged bNACHT11 of the indicated genotype. Representative image of n = 2 biological replicates.

**Fig. S6.**
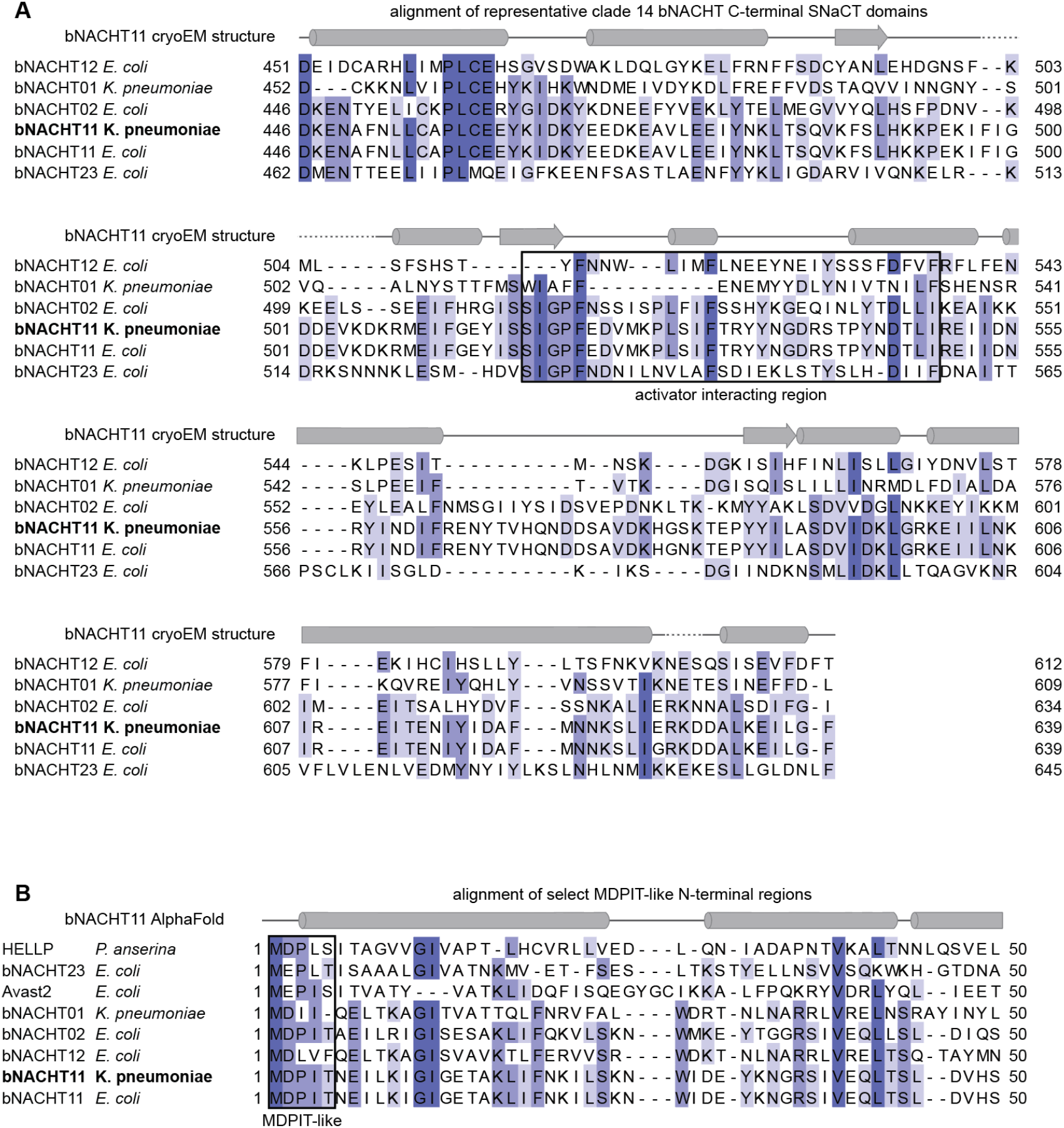
Protein alignments of bNACHT11 sensing and effector domains. **(A)** Alignment of SNaCT domains from representative clade 14 bNACHTs including bNACHT12, bNACHT01, bNACHT02, bNACHT11 *K. pneumoniae*, bNACHT11 *E. coli*, and bNACHT23. The secondary structure of bNACHT11 (PDB: 9ORF) is indicated above with alpha helices depicted as cylinders, β-sheets depicted as arrows, and unresolved regions of the structure depicted as a dashed line. A black box indicates the region with residues that appear to interact with phage protein activators. Amino acid residues are color coded based on percent identity such that darker colors indicate a higher degree percent identity in this alignment. **(B)** Alignment of N-terminal regions from bNACHT01, bNACHT02, bNACHT11 *K. pneumoniae*, bNACHT11 *E. coli*, bNACHT12, bNACHT23, HELLP, and Avast2. The secondary structure of bNACHT11 from an AlphaFold3 prediction is indicated above. A black box indicates MDPIT-like region.

**Fig S7.**
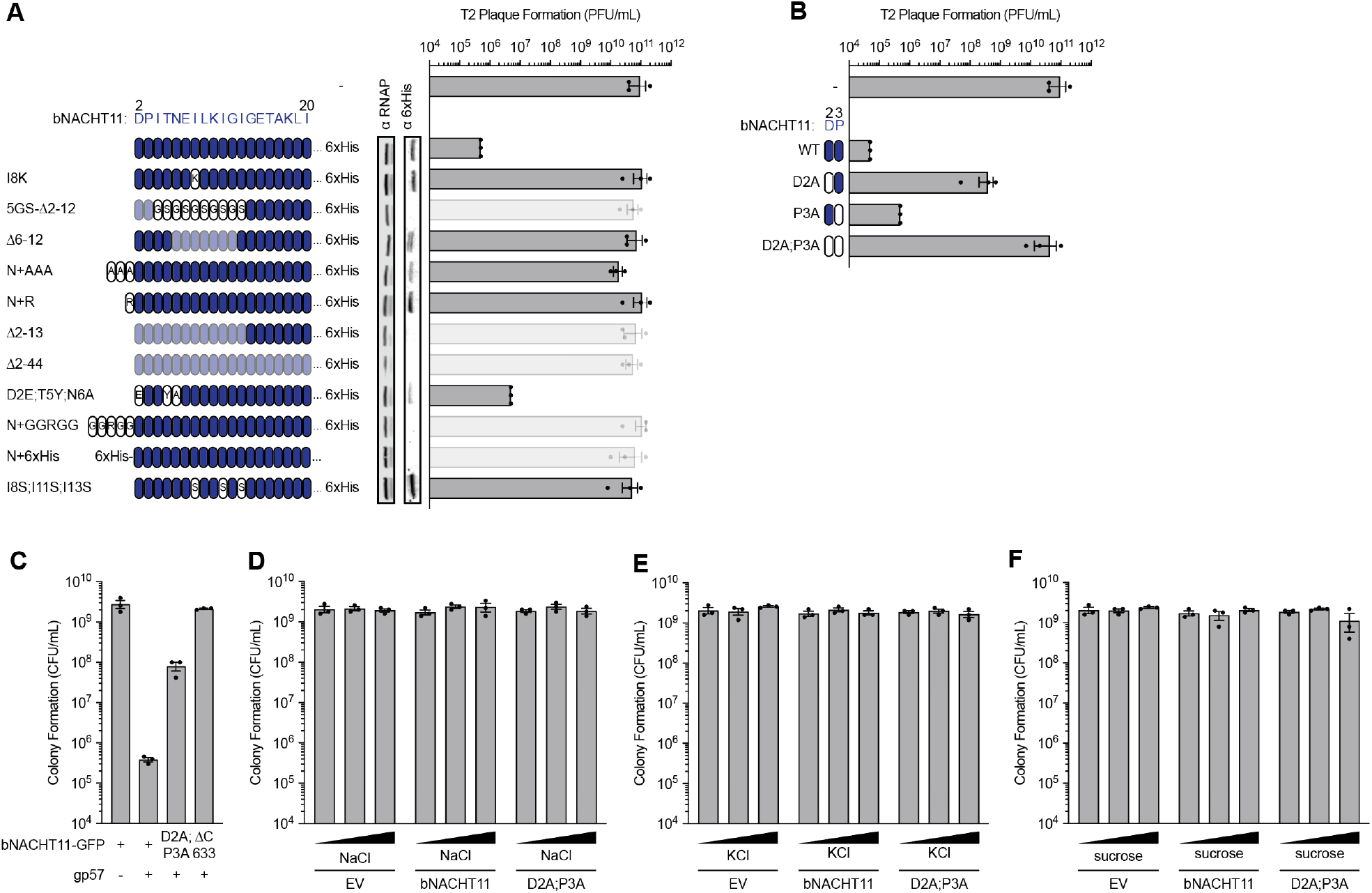
Additional mutations of bNACHT11 N-terminal extension, related to Figure 6. **(A)** Left: Cartoon depiction of deletions, additions, and substitutions made to the N-terminal extension of bNACHT11-6×His. WT residues are shown in navy blue, deleted residues in light blue, and substituted residues in white. Mutations resulting in no protein expression are indicated as bars with decreased opacity. Center: Western blot analysis of *E. coli* expressing 6×His-tagged bNACHT11 of the indicated genotype. Representative image of n = 2 biological replicates. Right: Efficiency of plating of phage T2 infecting *E. coli* expressing an empty vector (-) or the indicated genotype of bNACHT11-6×His. Data represent the mean ± SEM of n = 3 biological replicates, shown as individual points. **(B)** Left: Cartoon depiction of substitutions made to the N-terminal extension of bNACHT11. WT residues are shown in navy blue and residues mutated to alanine are white. Right: Efficiency of plating of phage T2 infecting *E. coli* expressing an empty vector (-) or the indicated genotype of bNACHT11. Data represent the mean ± SEM of n = 3 biological replicates, shown as individual points. **(C)** Quantification of colony formation of *E. coli* expressing gp57 on one plasmid and bNACHT11-GFP with the indicated genotype on a second plasmid. Data represent the mean ± SEM of n = 3 biological replicates, shown as individual points. **(D-F)** Quantification of colony formation of *E. coli* expressing an empty vector (EV), bNACHT11, or D2A; P3A, coexpressed with MBP. The expression of MBP is IPTG-inducible. Experiments were performed using 500 µM IPTG induction in LB media with the addition of (D) no NaCl (0 µM), normal NaCl (91 mM), high NaCl (300 mM), (E) no KCl (0 µM), normal KCl (91 mM), high KCl (300 mM), or (F) no sucrose (0 µM), normal sucrose (91 mM), high sucrose (300 mM). Data represent the mean ± SEM of n = 3 biological replicates, shown as individual points.

