## Supplemental Table 5 for "A bacterial NLR-related protein senses distinct phage triggers through a single interface"

**Table SX. Cryo-EM data collection, refinement and validation statistics**

|  | bNACHT11-gp57  (EMDB: EMD-70775)  (PDB 9ORF) |
| --- | --- |
| **Data collection and processing** |  |
| Magnification | 130,000 |
| Voltage (kV) | 300 |
| Electron exposure (e–/Å^2^) | 51 |
| Defocus range (μm) | -0.8 − -2.2 |
| Pixel size (Å) | 0.935 |
| Symmetry imposed | C7 |
| Initial particle images (no.) | 314,838 |
| Final particle images (no.) | 141,477 |
| Map resolution (Å)  FSC threshold | 2.8 − 5.67 |
| Map resolution range (Å) |  |
| **Refinement** |  |
| Initial model used (PDB code) | AlphaFold3 |
| Model resolution (Å)  FSC threshold | 2.5  0.143 |
| Map sharpening *B* factor (Å^2^) | 76.0 |
| Model composition  Non-hydrogen atoms  Protein residues  Ligands: ATP(Mg^2+^) | 36799  4473  7(7) |
| *B* factors (Å^2^)  Protein  Ligand | 132.74  76.89 |
| R.m.s. deviations  Bond lengths (Å)  Bond angles (°) | 0.002  0.412 |
| Validation  MolProbity score  Clashscore  Poor rotamers (%) | 1.51  4.07  0.51 |
| Ramachandran plot  Favored (%)  Allowed (%)  Disallowed (%) | 95.39  4.61  0 |
